# The Mediator complex regulates enhancer-promoter interactions

**DOI:** 10.1101/2022.06.15.496245

**Authors:** Shyam Ramasamy, Abrar Aljahani, Magdalena A. Karpinska, T. B. Ngoc Cao, J. Neos Cruz, A. Marieke Oudelaar

**Affiliations:** Max Planck Institute for Multidisciplinary Sciences, Göttingen, Germany

## Abstract

Enhancer-mediated gene activation generally requires physical proximity between enhancers and their target gene promoters. However, the molecular mechanisms by which interactions between enhancers and promoters are formed are not well understood. Here, we investigate the function of the Mediator complex in the regulation of enhancer-promoter interactions, by combining rapid protein depletion and high-resolution MNase-based chromosome conformation capture approaches. We show that depletion of Mediator leads to reduced enhancer-promoter interaction frequencies, which are associated with a strong decrease in gene expression. In addition, we find increased interactions between CTCF-binding sites upon Mediator depletion. These changes in chromatin architecture are associated with a re-distribution of the Cohesin complex on chromatin and a reduction in Cohesin occupancy specifically at enhancers. Our results indicate that enhancer-promoter interactions are dependent on an interplay between the Mediator and Cohesin complexes and provide new insights into the molecular mechanisms by which communication between enhancers and promoters is regulated.

## INTRODUCTION

Precise spatial and temporal patterns of gene expression in metazoans are regulated by enhancers, which are short non-coding DNA sequences that drive expression of their cognate gene promoters [1]. In mammals, enhancers can be located far upstream or downstream of the genes they control. To activate genes, enhancers interact with promoters in specific 3D chromatin structures [2]. Enhancer-mediated gene activation is therefore closely related to the three-dimensional organization of the genome [3]. However, the molecular mechanisms by which enhancer-promoter interactions are formed and enhancers drive gene expression remain incompletely understood.

Mammalian genomes are organized into compartments and Topologically Associating Domains (TADs). Compartments reflect separation of euchromatin and heterochromatin, whereas TADs represent relatively insulated regions of the genome, formed by loop extrusion [4]. In this process, ring-shaped Cohesin complexes translocate along chromatin and extrude progressively larger loops, until they are halted at CTCF-binding elements located at the boundaries of TADs [5]. Interacting enhancers and promoters are usually located in the same TAD [6]. Moreover, perturbations of TAD boundaries can cause ectopic enhancer-promoter interactions [7]. These observations suggest that loop extrusion could be involved in the regulation of enhancer-promoter communication and gene expression. Although it has been shown that depletion of components of the Cohesin complex does not lead to widespread mis-regulation of gene expression [8-10], Cohesin and its associated factors have been reported to be important for the regulation of cell type-specific genes [11-13]. In addition, it has recently been shown that depletion of Cohesin can cause weakening of enhancer-promoter interactions [14,15]. These observations suggest that Cohesin-mediated loop extrusion contributes to the formation of enhancer-promoter interactions. However, the molecular mechanism remains unclear. Furthermore, depletion of Cohesin causes a relatively subtle reduction in enhancer-promoter interaction strength [14]. This suggests that these interactions are not solely dependent on loop extrusion and that other mechanisms are involved in their formation.

Active enhancers and promoters are bound by transcription factors and coactivators, including the Mediator complex. Because the tail module of the Mediator complex interacts with the activation domains of transcription factors bound at enhancers and the head and middle modules interact with the pre-initiation complex (PIC) at gene promoters [16-18], it has been proposed that Mediator acts as a bridge between enhancers and promoters (reviewed in [19,20]). Initial studies based on knockdown of Mediator subunits over the course of several days provided evidence for this hypothesis [21-23]. However, since the Mediator complex has a central function in RNA Polymerase II (Pol II)-mediated transcription, its long-term perturbation causes secondary, confounding effects, which complicate the interpretation of these early studies.

To overcome these limitations, more recent studies have used rapid protein depletion strategies to investigate the function of the Mediator complex in gene regulation and genome organization [24,25]. These studies did not detect changes in chromatin architecture and enhancer-promoter interactions upon Mediator depletion, despite strongly reduced expression levels of enhancer-dependent genes. Based on these findings, it has been concluded that Mediator is dispensable for enhancer-promoter interactions and acts as a functional rather than an architectural bridge between enhancers and promoters [24,25].

A caveat of current studies of the role of Mediator in genome architecture is that enhancer-promoter interactions have been assessed with chromosome conformation capture (3C) methods at relatively low resolution [21-26]. It is therefore possible that changes in fine-scale genome architecture, including enhancer-promoter interactions, could not be reliably identified. For a better understanding of the function of the Mediator complex in genome regulation it is important to examine the impact of acute Mediator perturbations on chromatin architecture with high resolution and sensitivity.

Here, we overcome limitations of current studies and investigate the function of the Mediator complex by combining rapid protein depletion and high-resolution analysis of genome architecture using targeted MNase-based 3C approaches. We find that depletion of Mediator leads to a significant reduction of enhancer-promoter interactions. Interestingly, we also find that Mediator depletion causes increased interactions between CTCF-binding elements. We show that these changes in interaction patterns can be explained by a re-distribution of the Cohesin complex on chromatin and a specific loss of Cohesin occupancy at enhancers upon Mediator depletion. These results indicate that enhancer-promoter interactions are dependent on both Mediator and Cohesin and provide support for a model in which the Cohesin complex bridges and stabilizes interactions between enhancers and promoters bound by Mediator.

## RESULTS

### Mediator depletion causes changes in chromatin interaction patterns

Because the MED14 subunit acts as a central backbone that connects the Mediator head, middle and tail modules [19,20], its degradation disrupts the integrity of the Mediator complex [24,25]. We have therefore used an HCT-116 MED14-dTAG cell line [25] to study the function of the Mediator complex in genome regulation. Using immunoblotting, we have confirmed efficient MED14 depletion within two hours of treatment with a dTAG ligand (Supplementary Figure 1a).

Previous work has shown that Mediator depletion leads to strong downregulation of cell type-specific genes that are associated with super-enhancers [25] (Supplementary Figure 1b). Super-enhancers are stretches of clustered enhancers with high levels of Mediator that are thought to have a central role in driving high expression levels of key cell identity genes [27]. Previous studies could not detect changes in interactions between promoters and (super-)enhancers upon Mediator depletion [24-26]. However, these studies relied on genome-wide 3C approaches, such as Hi-C and HiChIP, with relatively low resolution (4-5 kb). It is therefore possible that small-scale changes in enhancer-promoter interactions could not be reliably detected.

To investigate changes in genome architecture upon Mediator depletion in more detail, we have used targeted 3C approaches, which are not limited by sequencing depth and can detect changes in genome structure at high resolution and with high sensitivity. We focused our analyses on 20 genes (Supplementary Figure 1b), which we selected based on the following criteria: (1) robust gene activity in HCT-116 cells; (2) significant down-regulation of gene expression upon Mediator depletion; (3) high Mediator occupancy at the gene promoter; (4) association with a super-enhancer. We initially used Capture-C [28,29], a targeted 3C method based on DpnII digestion, to evaluate changes in chromatin interactions with the promoters of these genes. Capture-C interaction profiles display interaction frequencies with selected viewpoints per DpnII restriction fragment and therefore have an average resolution of ∼250 bp. By comparing Capture-C data generated in HCT-116 MED14-dTAG cells treated with DMSO or dTAG ligand, we find that Mediator depletion leads to subtle changes in the interaction patterns of the selected gene promoters (Figure 1, Supplementary Figure 2). Unexpectedly, we observe patterns of both decreased and increased interactions.

**Figure 1.**
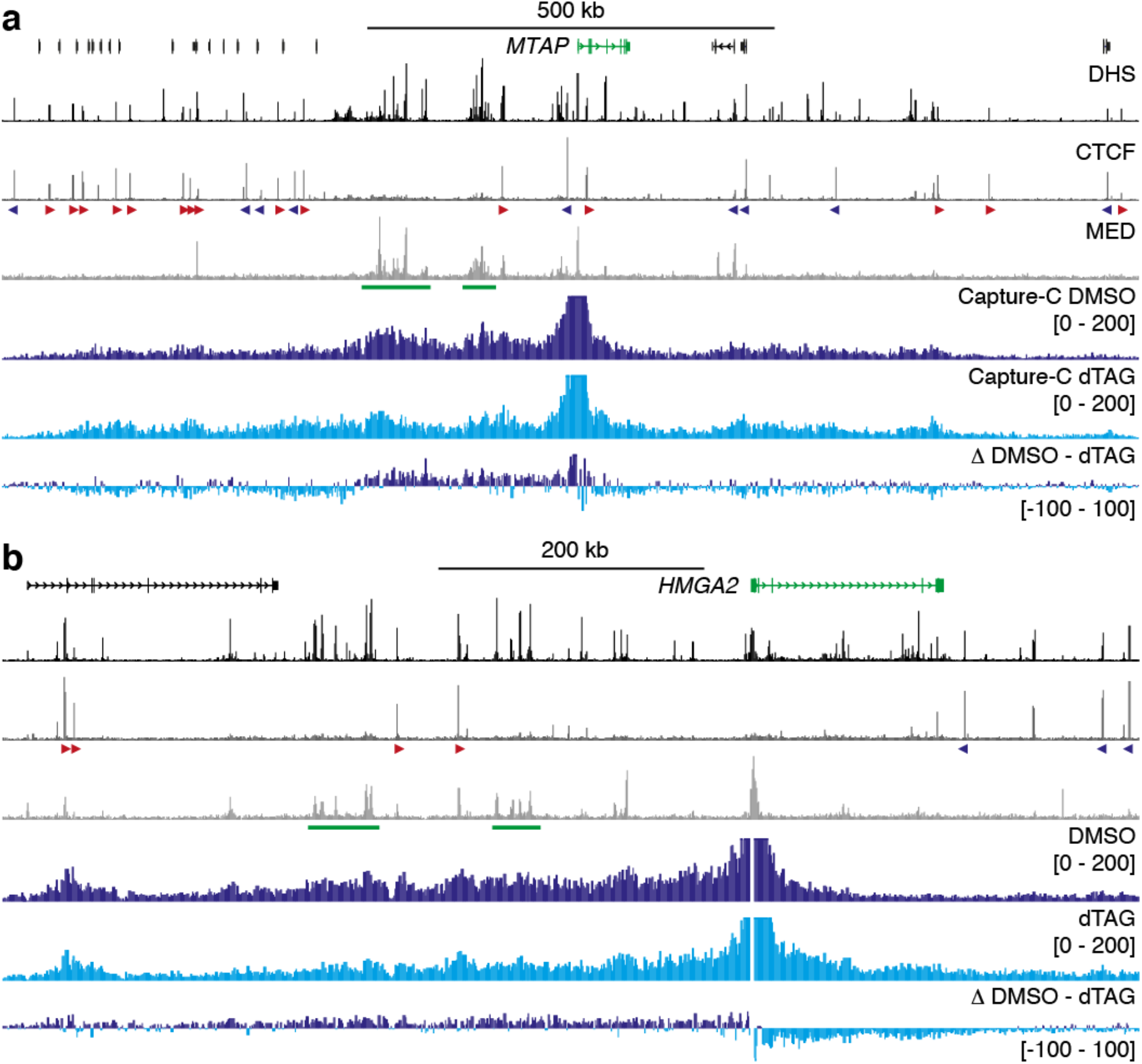
Changes in chromatin interactions upon Mediator depletion. **a**. Normalized Capture-C interaction profiles from the viewpoint of the *MTAP* promoter in HCT-116 MED14-dTAG cells treated for two hours with DMSO (dark blue) or dTAG ligand (light blue). Gene annotation, DNase hypersensitive sites (DHS), and ChIP-seq data for CTCF and MED26 are shown above. Enhancers of interest are highlighted in green below the MED26 profiles and orientations of CTCF motifs are indicated with arrowheads (forward orientation in red; reverse orientation in blue). The axes of the DHS and ChIP-seq profiles are scaled to signal and have the following ranges: DHS = 0–186; CTCF = 0–112; MED26 = 0–62. Coordinates (hg38): chr9:21,096,000-22,491,000. **b**. Data as described in **a** for the *HMGA2* locus. The axes of the DHS and ChIP-seq profiles are scaled to signal and have the following ranges: DHS = 0–181; CTCF = 0–93; MED26 = 0–66. Coordinates (hg38): chr12:65,260,000-66,115,000.

For example, in the *MTAP* locus, the Capture-C data show reduced interactions in the region upstream in which two super-enhancers are located, and a tendency for increased interactions with the regions further upstream and downstream (Figure 1a). In the region containing the *HMGA2* oncogene, interactions with the region upstream, which contains two super-enhancers, are weakened, whereas downstream interactions are increased (Figure 1b). We observe similar patterns of both decreased and increased interactions in other loci we investigated (Supplementary Figure 2).

### Depletion of Mediator leads to reduced enhancer-promoter interactions

To examine the broad changes in the Capture-C interaction profiles in further detail, we performed Micro-Capture-C (MCC) experiments [30] in DMSO- and dTAG-treated HCT-116 MED14-dTAG cells, using viewpoints targeting the same set of gene promoters. Compared to Capture-C, MCC has the advantage that it uses MNase instead of DpnII for chromatin digestion. The resolution of MCC is therefore not limited by the distribution of DpnII cut sites across the genome, enabling analysis at base-pair resolution [30]. The MCC data resolve the broad interaction patterns in the Capture-C data and clearly show that Mediator depletion leads to reduced interactions between gene promoters and Mediator-bound enhancer regions (Figure 2, Supplementary Figure 3).

**Figure 2.**
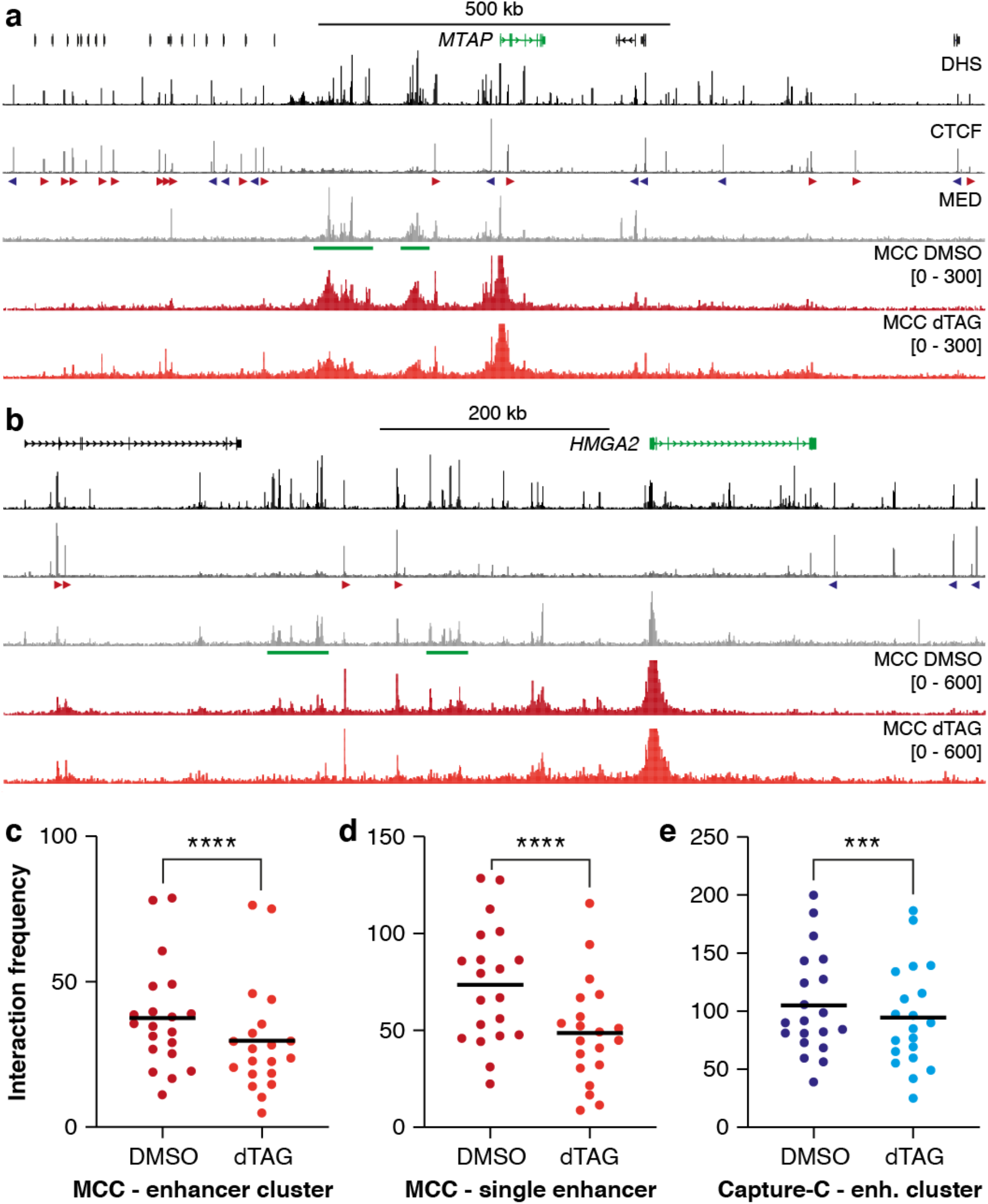
Depletion of Mediator leads to decreased enhancer-promoter interactions and increased interactions with CTCF-binding sites. **a**. Normalized Micro-Capture-C interaction profiles from the viewpoint of the *MTAP* promoter in HCT-116 MED14-dTAG cells treated for two hours with DMSO (dark red) or dTAG ligand (light red). Gene annotation, DNase hypersensitive sites (DHS), and ChIP-seq data for CTCF and MED26 are shown above. Enhancers of interest are highlighted in green below the MED26 profiles and orientations of CTCF motifs are indicated with arrowheads (forward orientation in red; reverse orientation in blue). The axes of the DHS and ChIP-seq profiles are scaled to signal and have the following ranges: DHS = 0–186; CTCF = 0–112; MED26 = 0–62. Coordinates (hg38): chr9:21,096,000-22,491,000. **b**. Data as described in **a** for the *HMGA2* locus. The axes of the DHS and ChIP-seq profiles are scaled to signal and have the following ranges: DHS = 0–181; CTCF = 0–93; MED26 = 0–66. Coordinates (hg38): chr12:65,260,000-66,115,000. **c**. Quantification of interaction frequencies between gene promoters and enhancer clusters (average size: 58 kb) extracted from MCC data in 20 loci. **** P < 0.0001 (paired t-test). **d**. Quantification of interaction frequencies between gene promoters and individual enhancers (average size: 2.7 kb) extracted from MCC data in 20 loci. **** P < 0.0001 (paired t-test). **e**. Quantification of interaction frequencies between gene promoters and enhancer clusters (average size: 58 kb) extracted from the Capture-C data presented in Figure 1 in 20 loci. *** P = 0.0002 (paired t-test).

By quantifying the MCC interactions between gene promoters and clusters of Mediator-bound enhancers, we find that Mediator depletion leads to an average reduction in interaction frequency of 22% in the 20 regions we focused on (Figure 2c). The reduction in interaction frequency between the promoters and a narrow region covering the largest Mediator peak within these broad clusters is 34% (Figure 2d). These changes are associated with an average 7.5-fold decrease in gene expression (Supplementary Figure 1b). Of note, the reduction in interactions between promoters and enhancer clusters in Capture-C data is 9% (Figure 2e). Although consistent with the MCC data, this comparison highlights the need for analyses with sufficient resolution and sensitivity to robustly detect changes in enhancer-promoter interactions.

### Interactions with CTCF-binding sites are increased in absence of Mediator

The MCC data do not only identify precise reductions in interactions with enhancers, but also uncover very punctate increased interactions in absence of Mediator. Strikingly, these increased interactions all overlap with CTCF-binding sites. For example, in the CTCF-dense *MTAP* locus, we see strong increases in interactions formed with CTCF-binding sites in the region upstream of the super-enhancers and downstream of the gene promoter (Figure 2a). Notably, the interacting CTCF-binding sites upstream are all in a forward orientation, whereas the interacting CTCF -binding sites downstream are all in a reverse orientation.

We observe a similar pattern of increased interactions with convergently orientated CTCF-binding sites in the *MYC* locus after Mediator depletion (Supplementary Figure 3c). In the *HMGA2, ERRF1, KRT19* and *ITPRID2* loci, which contain fewer CTCF-binding sites, the patterns are a bit more subtle, but also clearly present (Figure 2b, Supplementary Figure 3a,b,d).

### Mediator depletion leads to a redistribution of intra-TAD interactions

The MCC data show clear and precise changes in chromatin interactions in absence of the Mediator complex. However, since the MCC viewpoints are very narrow and focused on gene promoters, it remains unclear how large-scale 3D genome architecture is changed and how interactions between other c*is*-regulatory elements are impacted by Mediator depletion. We therefore used the Tiled-MCC approach, in which MCC library preparation is combined with an enrichment strategy based on capture oligonucleotides tiled across large genomic regions of interest [14], to investigate changes in genome architecture in DMSO- and dTAG-treated HCT-116 MED14-dTAG cells in a broader context. We focused on the *MYC* locus (3.3 Mb; Figure 3), *MTAP* locus (1.55 Mb; Supplementary Figure 4), *HMGA2* locus (990 kb; Supplementary Figure 5) and *ITPRID2* locus (900 kb; Supplementary Figure 6).

**Figure 3.**
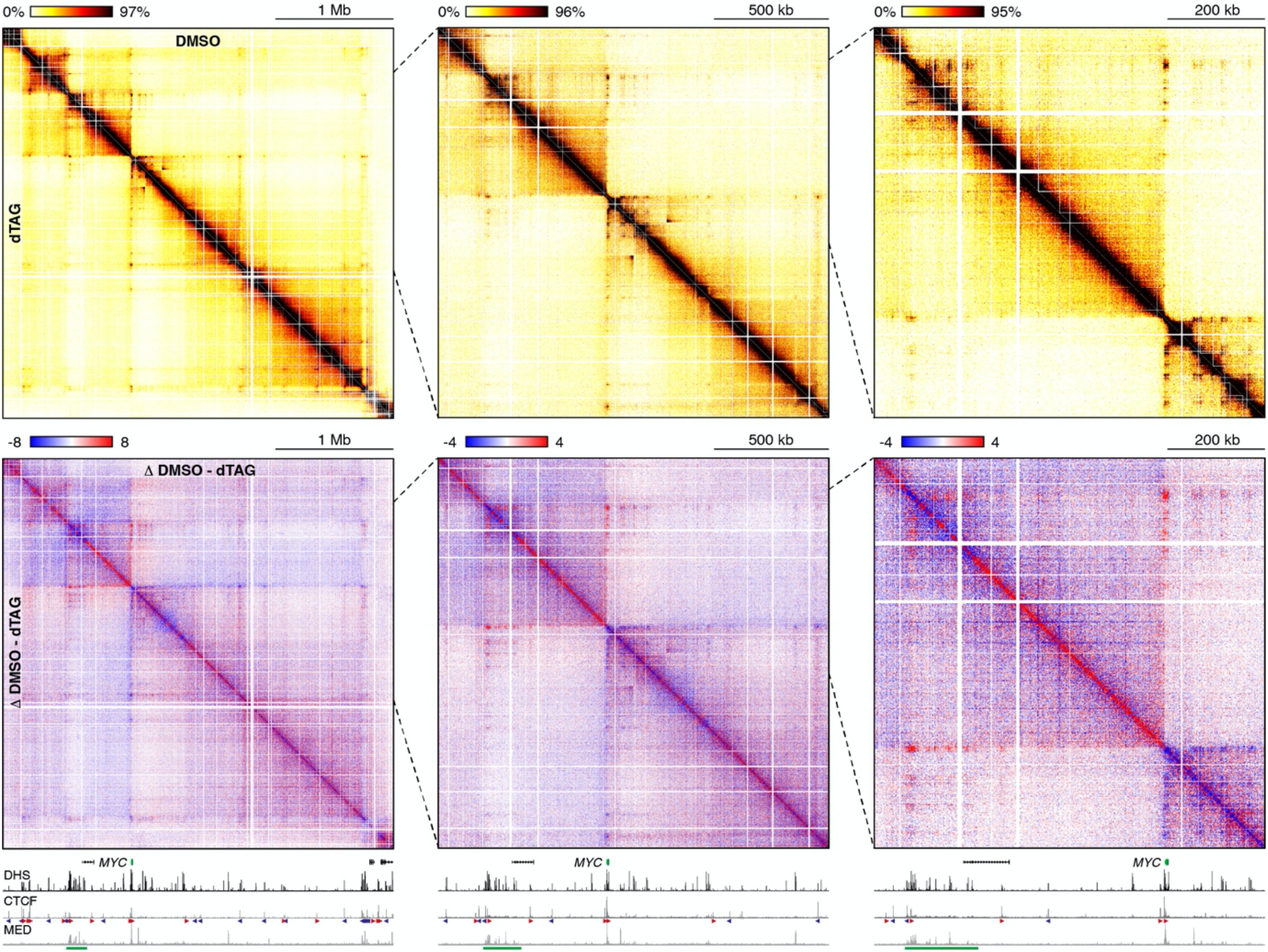
Mediator depletion results in subtle changes in large-scale genome organization. Tiled-MCC contact matrices of the *MYC* locus in HCT-116 MED14-dTAG cells treated for two hours with DMSO (top-right) or dTAG ligand (bottom-left). The right matrices show two different zoomed views, of which the areas are indicated by the black dashed lines. Differential contact matrices, in which interactions enriched in DMSO-treated cells are shown in red and interactions enriched in dTAG-treated cells are shown in blue, are displayed below. Gene annotation, DNase hypersensitive sites (DHS), and ChIP-seq data for CTCF and MED26 are shown at the bottom. Enhancers of interest are highlighted in green below the MED26 profiles and orientations of CTCF motifs are indicated with arrowheads (forward orientation in red; reverse orientation in blue). The axes of the DHS and ChIP-seq profiles are scaled to signal and have the following ranges: DHS = 0–188; CTCF = 0–188; MED26 = 0–64. Coordinates (hg38): chr8:126,650,000-129,950,000

In line with previous studies that examined changes in genome architecture using Hi-C and Hi-ChIP [24-26], we do not detect drastic changes in large-scale genome organization after Mediator depletion. We find that TAD organization is preserved, without any shifts in the location of boundaries. However, we do find subtle changes in interaction patterns within TADs. In line with the Capture-C and MCC data, we observe that enhancer-promoter interactions are reduced in absence of Mediator. In addition, we detect strengthening of interactions anchored at CTCF-binding sites. As a result, we see subtle increases in “looping” between the CTCF-bound anchors of TADs and sub-TADs.

### Cohesin occupancy patterns are altered after Mediator depletion

It has been shown that CTCF and Cohesin co-localize and that interactions between CTCF-binding sites are formed via loop extrusion by the Cohesin complex [31-34]. Notably, Cohesin also co-localizes with Mediator and co-immunoprecipitation experiments have suggested that these complexes interact [21,22]. However, a functional link between Mediator and Cohesin has not been identified.

Since our data show that depletion of Mediator causes a decrease in enhancer-promoter interactions and an increase in CTCF-mediated interactions, we hypothesized that these altered interaction patterns could be explained by changes in the distribution of the Cohesin complex on chromatin. To test this, we mapped Cohesin occupancy using Cleavage Under Targets and Tagmentation (CUT&Tag [35]) in DMSO- and dTAG-treated HCT-116 MED14-dTAG cells (Figure 4, Supplementary Figure 7). These data show clear changes in Cohesin occupancy upon Mediator depletion. For example, in the *ERRFI1* locus we observe a reduction in Cohesin levels at the Mediator-bound elements of the super-enhancer (Figure 4a), which is associated with a loss of interactions between the super-enhancer and the *ERRFI1* promoter (Supplementary Figure 3a). However, Cohesin occupancy at the close-by CTCF-binding sites is not affected by Mediator depletion. At the *KRT19* locus we also observe weaker Cohesin occupancy at the Mediator-bound elements within the super-enhancer (Figure 4b, Supplementary Figure 3b)). This super-enhancer also contains a CTCF-binding site, at which Cohesin levels remain stable in absence of Mediator. We find similar patterns in the *MTAP, HMGA2, MYC* and *ITPRID2* loci (Supplementary Figure 7). Genome-wide quantification of Cohesin occupancy at Mediator-bound enhancers and CTCF-binding sites shows a clear reduction in Cohesin levels at enhancers and stable occupancy at CTCF-binding sites in absence of Mediator (Figure 4c,d). These results show that the distribution of Cohesin is altered in absence of Mediator and indicate that Mediator contributes to the stabilization of Cohesin at enhancer elements.

**Figure 4.**
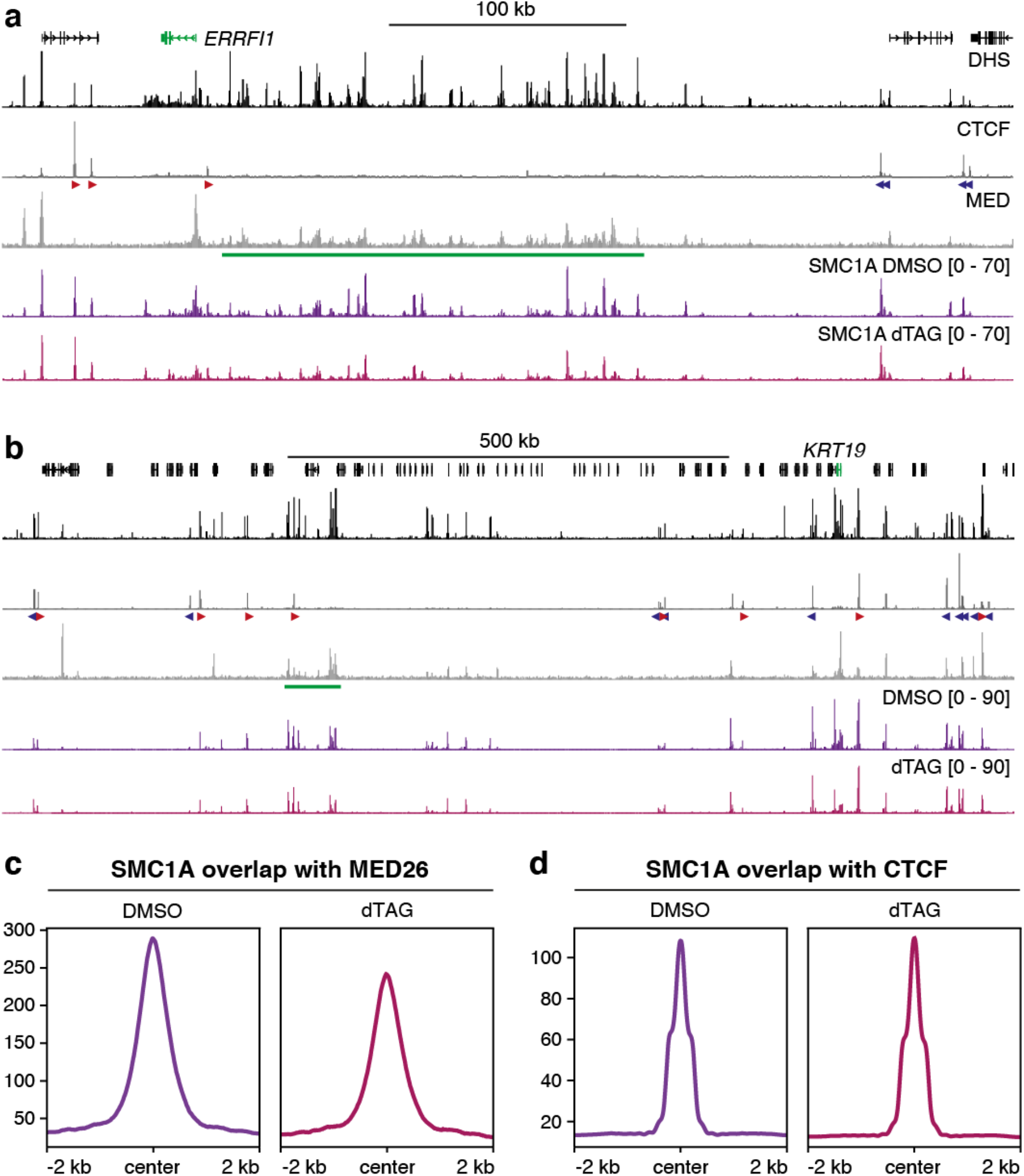
Cohesin occupancy at enhancers is reduced in absence of Mediator. **a**. CUT&Tag data for the Cohesin subunit SMC1A in the *ERRFI1* locus in HCT-116 MED14-dTAG cells treated for two hours with DMSO (dark purple) or dTAG ligand (light purple). Gene annotation, DNase hypersensitive sites (DHS), and ChIP-seq data for CTCF and MED26 are shown above. Enhancers of interest are highlighted in green below the MED26 profiles and orientations of CTCF motifs are indicated with arrowheads (forward orientation in red; reverse orientation in blue). The axes of the DHS and ChIP-seq profiles are scaled to signal and have the following ranges: DHS = 0–170; CTCF = 0–253; MED26 = 0–56. Coordinates (hg38): chr1:7,945,000-8,370,000. **b**. Data as described in **a** for the *KRT19* locus. The axes of the DHS and ChIP-seq profiles are scaled to signal and have the following ranges: DHS = 0–180; CTCF = 0–301; MED26 = 0–77. Coordinates (hg38): chr17:40,580,000-41,725,000. **c**. Meta-analysis of SMC1A peaks overlapping with MED26 peaks in HCT-116 MED14-dTAG cells treated for two hours with DMSO (dark purple) or dTAG ligand (light purple). **d**. Meta-analysis of SMC1A peaks overlapping with CTCF peaks in HCT-116 MED14-dTAG cells treated for two hours with DMSO (dark purple) or dTAG ligand (light purple).

### Mediator depletion results in changes in nano-scale interaction patterns

To further analyze the impact of Mediator depletion on chromatin architecture, we leveraged the ability of Tiled-MCC to directly identify ligation junctions and resolve localized nano-scale interaction patterns [14]. We focused our analyses on ligation junctions in regions containing super-enhancers, genes and boundary elements in the *MYC, MTAP, HMGA2*, and *ITPRID2* loci (Figure 5, Supplementary Figures 8-10).

Within the *MYC* super-enhancer, we observe enriched interactions between the individual elements of the super-enhancer (Figure 5, left matrix). Upon depletion of Mediator, these interactions are decreased. We observe similar patterns in the *MTAP, HMGA2* and *ITPRID2* loci (Supplementary Figures 8-10). In the *MTAP* locus, we could also resolve the interactions between the gene promoter and a nearby enhancer. In absence of Mediator, these interactions are reduced (Supplementary Figure 8). These results show that interactions between active enhancer and promoter elements across very small distances are dependent on Mediator.

**Figure 5.**
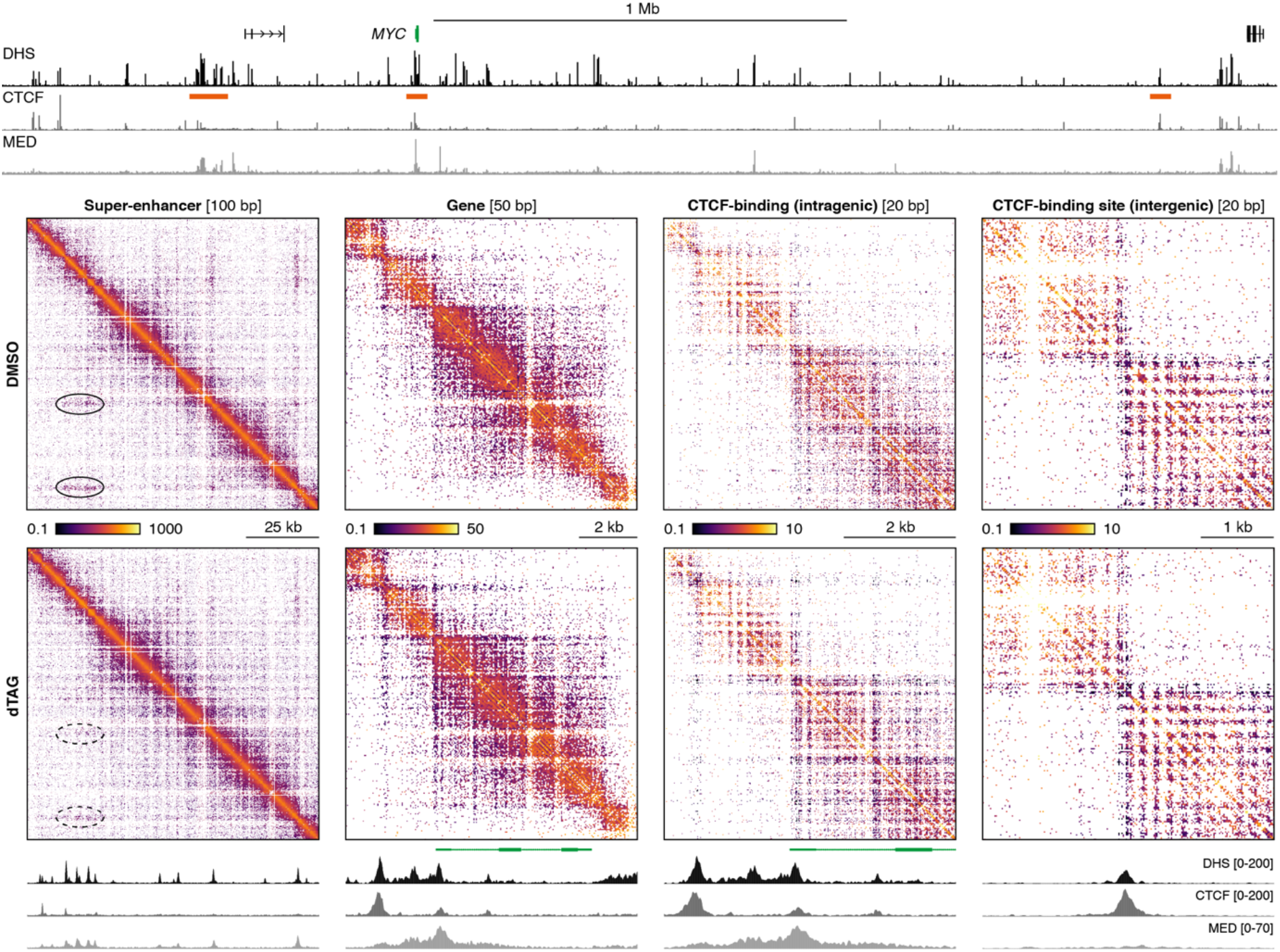
Depletion of Mediator leads to changes in nano-scale genome organization. Tiled-MCC ligation junctions in the *MYC* locus in HCT-116 MED14-dTAG cells treated for two hours with DMSO (top) or dTAG ligand (bottom), displayed in localized contact matrices at high resolution. Gene annotation, DNase hypersensitive sites (DHS), and ChIP-seq data for CTCF and MED26 for the extended and localized *MYC* locus are shown above and below the matrices, respectively. The regions covered in the contact matrices are highlighted with orange bars (not drawn to scale) below the top DHS profile and show a super-enhancer, a gene, an intragenic CTCF-binding site, and an intergenic CTCF-binding site, at the indicated resolution. Note that the intragenic CTCF-binding site in the third matrix is contained within the gene displayed in the second matrix. The axes of the DHS and ChIP-seq profiles at the bottom are fixed as indicated; the profiles at the top are scaled to signal with the following ranges: DHS = 0–188; CTCF = 0–188; MED26 = 0–64.

It has previously been shown that regions containing CTCF-binding sites form characteristic architectural patterns, in which phased nucleosomes surrounding the CTCF motif form a grid-like structure, which is associated with strong insulation between the regions upstream and downstream of the CTCF-binding site [14,36]. We observe these patterns at the intergenic CTCF-binding sites in the loci we investigated and do not see any changes upon depletion of Mediator (Figure 5, right matrix, Supplementary Figures 8-10).

At the level of individual genes, we observe domain-like structures extending across the gene body (Figure 5, Supplementary Figures 8-10). Interestingly, we observe that depletion of Mediator results in the appearance of specific structures within the *MYC* gene, which are centered around hypersensitive and CTCF-bound elements (Figure 5, left middle matrix). Zooming in to this region at higher resolution (Figure 5, right middle matrix) resolves a structure that is reminiscent of intergenic CTCF-binding sites at the CTCF-bound region within the *MYC* gene body in absence of Mediator. This suggests that high transcriptional activity in the presence of Mediator leads to a disruption of the specific nucleosome structures that are normally formed around CTCF-binding sites. We observe similar patterns at the CTCF-binding sites contained within the *MTAP* and *ITPRID2* gene bodies upon depletion of Mediator (Supplementary Figures 8,10). The observation that specific higher-order nucleosome structures can only form in absence of high transcriptional activity is consistent with experiments in yeast that have shown that the transcriptional machinery disrupts regular nucleosome spacing [37].

### BET proteins do not compensate for Mediator loss

Our data show that both short- and long-range interactions between enhancers and promoters are dependent on Mediator. However, we find that enhancer-promoter interactions are not completely abolished in absence of Mediator. This suggests that other factors are involved in mediating enhancer-promoter interactions and possibly compensate for the absence of Mediator. It has recently been shown that BRD4 plays a role in genome organization and stabilizes Cohesin on chromatin [38]. Although it has been shown that inhibition of BET proteins alone does not lead to reduced enhancer-promoter interactions [39], we wondered whether Mediator and BET proteins might have (partly) redundant roles in enhancer-promoter interactions. We therefore investigated the impact of combined Mediator depletion and chemical BET inhibition on enhancer-promoter interactions with Capture-C (Supplementary Figure 11). However, we did not find consistent additional effects on enhancer-promoter interactions after combined Mediator depletion and BET inhibition compared to depletion of Mediator alone. This suggests that enhancer-promoter interactions result from a more complex interplay between a multitude of regulatory factors.

## DISCUSSION

In this study, we have investigated the function of the Mediator complex in the regulation of chromatin architecture and enhancer-promoter interactions. To overcome limitations of existing studies [21-26], we have combined rapid depletion of Mediator using dTAG technology and analysis of genome architecture at very high resolution with targeted MNase-based 3C approaches. This strategy has enabled us to demonstrate that depletion of Mediator leads to a significant reduction in enhancer-promoter interactions (Figure 6).

**Figure 6.**
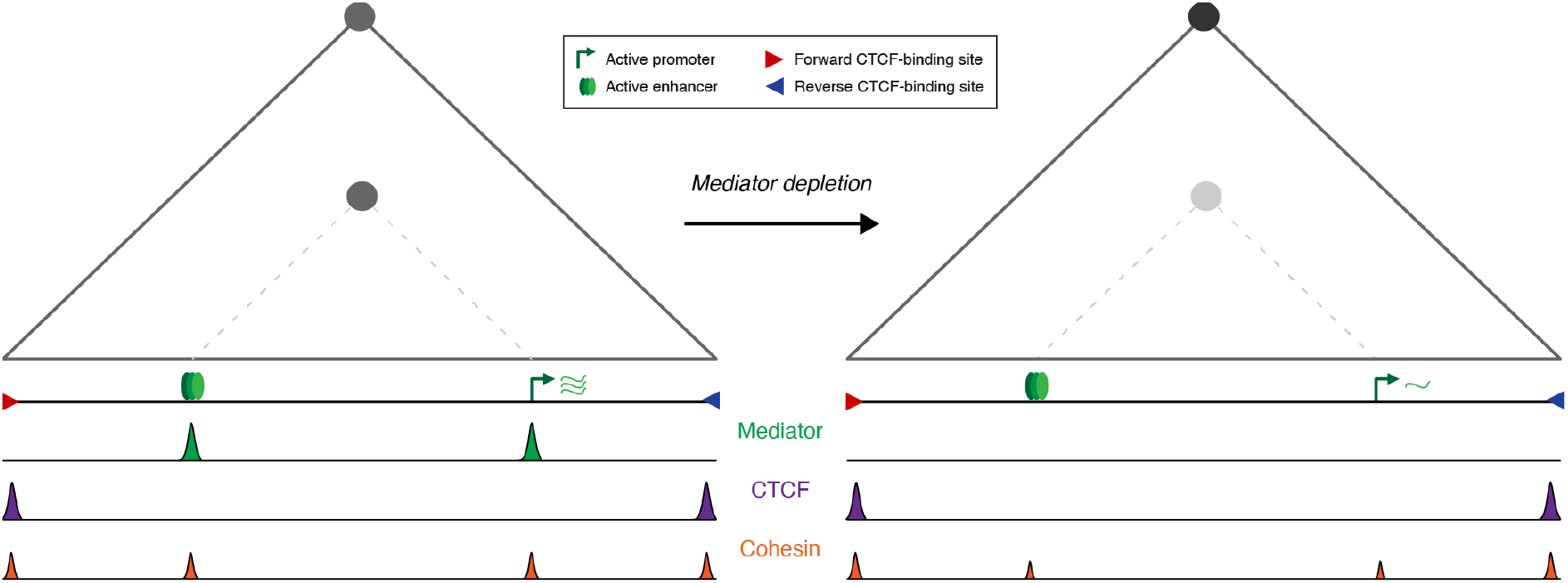
Graphical summary. The left panel shows a schematic TAD (grey triangle), interactions between the CTCF-bindings sites located at its boundaries (grey circle at the TAD apex), and enhancer-promoter interactions (grey circle at the intersection between the enhancer and promoter, as indicated with a dashed line). Upon Mediator depletion, Cohesin occupancy at the enhancer is reduced and enhancer-promoter interactions are weakened. However, the TAD structure remains intact and the interactions between the CTCF-binding sites at the TAD boundaries are increased.

We have focused our analyses on 20 gene loci containing strong super-enhancers and find an average decrease in interaction strength between promoters and Mediator-bound enhancer elements in these regions of ∼34%. This reduction in enhancer-promoter interactions is associated with an average ∼7.5-fold downregulation of expression of the genes we investigated. The relatively small effect on interaction frequency in comparison with gene activity is in agreement with recent studies, which have shown that the relationship between enhancer-promoter interaction frequency and transcriptional output is not linear and that small changes in genome architecture can have a large impact on gene activity levels [40,41].

In the context of Mediator depletion, there are several possible explanations for these observations. We have focused our analyses on genes regulated by super-enhancers, which are composed of many individual elements. For example, the *MTAP* gene is regulated by two super-enhancers, which contain more than twenty individual active elements. The additive and potentially synergistic impact of reduced interactions of each of these elements could cumulatively cause large changes in gene expression levels. In addition, the Mediator complex plays a central role in the regulation of gene expression and is thought to act at several stages of the transcription cycle [19,20]. It is therefore likely that the large decrease in transcriptional output upon Mediator depletion is not only related to weaker enhancer-promoter interactions, but also to the loss of the general function of Mediator in initiation, re-initiation and elongation. Moreover, it is thought that the function of the Mediator complex in gene regulation is (partly) dependent on the formation of nuclear condensates [42-46]. In agreement with this model, it has been shown that MED14 depletion leads to dissolved Pol II clusters [25]. It is possible that the reduced interactions between enhancers and promoters in absence of Mediator are not sufficient to establish the required concentrations of transcription factors, coactivators, and Pol II for the formation of nuclear condensates in which transcription can be efficiently initiated.

Our data indicate that Mediator’s role in enhancer-promoter interactions is (partly) dependent on Cohesin. Although it has previously been shown that Mediator co-localizes with Cohesin [21,22], the functional relationship between these complexes has thus far been unclear. Our data show that Cohesin levels at enhancers are reduced in absence of Mediator, which indicates that Mediator stabilizes Cohesin on chromatin. This suggests that Cohesin and Mediator cooperate in the formation of enhancer-promoter interactions and provides support for a model in which extruding Cohesin molecules are stalled at Mediator-bound enhancers and promoters and thereby bridge interactions between these elements. These findings indicate that Cohesin extrusion trajectories are dependent on multiple regulatory proteins and that these factors cooperate in the formation of specific three-dimensional chromatin structures in which gene expression is regulated [47,48].

The high resolution of our data has enabled us to visualize the effects of Mediator depletion on nano-scale genome organization. We find that interactions between the individual elements within super-enhancers and interactions between enhancers and promoters across very small distances are dependent on Mediator. Of note, it has previously been shown that Cohesin depletion leads to reduction of enhancer-promoter interactions across medium and large genomic distances (> ∼10 kb), but that Cohesin is not involved in regulating short-range enhancer-promoter interactions and interactions within enhancer clusters [14]. This suggests that Cohesin has a facilitating role in longer-range enhancer-promoter interactions and that Mediator can function independently on smaller scales. At the level of nano-scale genome organization, we also observe that reduced transcription following Mediator depletion leads to the formation of specific chromatin structures within gene bodies, particularly around CTCF-binding elements. This shows that higher-order nucleosome structures at CTCF-binding sites are incompatible with high levels of transcription.

With the exception of a subtle increase in the strength of TAD and sub-TAD boundaries, we do not observe large-scale changes in genome architecture upon Mediator depletion. This is consistent with previous reports, in which the impact of Mediator depletion has been investigated with lower resolution approaches such as Hi-C and Hi-ChIP [24-26]. Based on knock-out of the Mediator-CDK module, it has recently been suggested that the Mediator complex is involved in the regulation of heterochromatin domains and genome compartmentalization [49]. We do not observe clear changes in compartmentalization after two hours of Mediator depletion, but it is likely that changes in compartmentalization would require more time to manifest [50-52].

Although our data clearly show that enhancer-promoter interactions are dependent on Mediator, we do not observe a complete loss of interactions in absence of Mediator. This suggests that additional proteins and mechanisms play a role in mediating enhancer-promoter interactions. We find that the remaining interactions in absence of Mediator are not dependent on BET proteins. However, many other regulatory factors, such as tissue-specific transcription factors [53-55] and more widely expressed transcription factors such as LDB1 [56-59] and YY1 [60,61] have been implicated in enhancer-promoter interactions. It is likely that the regulation of enhancer-promoter interactions is dependent on a complex interplay between multiple regulatory proteins, which might act in a (partly) redundant manner to ensure the formation of robust enhancer-promoter interactions. In line with biochemical and structural evidence [19,20], our data show that the Mediator complex is one of the factors with an important role in regulating enhancer-promoter communication and gene expression, by acting as both a functional and an architectural bridge between enhancers and promoters.

## METHODS

### Cell culture

Wild type and MED14-dTAG human colorectal carcinoma HCT-116 cells [25] were grown in RPMI 1640 medium (Gibco, 21875034) supplemented with 10% FBS (Gibco, 10270106) and 1X Penicillin-Streptomycin (Gibco, 15140122) at 37°C and 5% CO^2^. Cells were passaged once every 2-3 days by trypsinization (Gibco, 25300054) upon reaching ∼70-80% confluency. For MED14 depletion, HCT-116 MED14-dTAG cells were seeded in 175 cm^2^ culture flasks. At 80% confluency, the cells were replenished with fresh culture medium containing 0.5 mM dTAG^v^-1 (Tocris, 6914) and treated for 2 h. For co-inhibition of BET proteins, MED14 depletion was combined with treatment with I-BET 151 dihydrochloride (Tocris, 4650), a BET bromodomain inhibitor, for 90 min. HCT-116 MED14-dTAG cells treated with 0.1% DMSO served as a control in all experiments.

### Immunoblotting

For validation of MED14 depletion, HCT-116 MED14-dTAG cells were treated with dTAG^v^-1 for 0.5, 1, 2, 4, 6 and 8 h. Following treatment, the cells were trypsinized and pelleted. The cell pellets were washed once with PBS and lysed in radioimmunoprecipitation assay (RIPA) lysis and extraction buffer (Thermo Scientific, 89900) supplemented with 250 U/mL Benzonase (Sigma-Aldrich, E1014) and a protease inhibitor cocktail containing Leupeptin (Carl Roth, CN33.4), PMSF (Carl Roth, 6367.3), Pepstatin A (Carl Roth, 2936.3) and Benzamide hydrochloride (Acros Organics, E1014) for 1 h at 4°C on a rotator. Lysates were cleared by centrifugation at maximum speed for 15 min at 4°C. Protein concentration was measured using the Bio-Rad Protein Assay (Bio-Rad, 5000006). For each sample, 20 µg of protein lysate was mixed with 4X LDS sample buffer (Invitrogen, NP0007), supplemented with 50 mM DTT (Carl Roth, 6908.3) and denatured for 5 min at 95°C. Proteins were separated on a NuPAGE 4-12% Bis-Tris gel (Invitrogen, NP0321) and blotted to a PVDF membrane. The membrane was blocked with 5% milk (Carl Roth, T145.2) in 1X PBS containing 0.05% Tween-20 (PBST) for 1 h at room temperature and was cut into two parts to detect higher and lower molecular weight target proteins separately. Cut membranes were incubated with primary antibodies (MED14-HA: 1:1000, HA-Tag (C29F4) antibody, rabbit monoclonal, Cell Signaling Technology, 3724S; GAPDH loading control: 1:5000, anti-GAPDH antibody (6C5), mouse monoclonal, Abcam, ab8245) in 5% milk at 4°C overnight, followed by 3 washes with PBST for 10 min each and incubation with HRP-labeled secondary antibodies (MED14-HA: 1:3000, Goat Anti-Rabbit IgG H&L (HRP), Abcam, ab205718; GAPDH loading control: 1:3000, Goat Anti-Mouse IgG H&L (HRP), Abcam, ab205719) for 1 h at room temperature. The blots were washed 3 times with PBST again, developed and imaged using an INTAS ChemoCam Imager HR.

### Capture-C

Capture-C was performed as described previously [62,63] for three biological replicates per experimental condition. Briefly, 10 million cells per biological replicate were crosslinked, followed by cell lysis. 3C libraries were generated by DpnII digestion and subsequent proximity ligation. After decrosslinking and DNA extraction, the resulting 3C libraries were sonicated to a fragment size of ∼200 bp and indexed with Illumina sequencing adapters, using Herculase II polymerase (Agilent, 600677) for library amplification. To boost library complexity, indexing was performed in two parallel reactions for each sample. Biotinylated oligonucleotides (70 nt) were designed using a python-based oligo tool [64] (https://oligo.readthedocs.io/en/latest/) and ordered from IDT as xGen Lockdown Probe Pools. The oligonucleotides were used for enrichment of the libraries in two consecutive rounds of hybridization, biotin-streptavidin bead pulldown (Invitrogen, 65306), bead washes and PCR amplification (KAPA HyperCapture Reagent Kit, Roche, 09075828001). The final libraries were assessed on a fragment analyzer using the High Sensitivity NGS Analysis Kit (Agilent, DNS-474-FR) and sequenced using the NextSeq550 Illumina platform (75-bp paired-end reads). Data analysis was performed using the CapCruncher pipeline [62] (https://github.com/sims-lab/CapCruncher).

### Micro-Capture-C

Micro-Capture-C (MCC) was performed as described previously [30] for three biological replicates per experimental condition. Briefly, multiple aliquots of 10 million cells per biological replicate were crosslinked and permeabilized with 1% digitonin (Sigma-Aldrich, D141). For each replicate, the permeabilized cells were pelleted, resuspended in nuclease-free water, and split into three digestion reactions. MCC libraries were generated by digesting the chromatin in low Ca^2+^ MNase buffer (10 mM Tris-HCl pH7.5, 10 mM CaCl^2^) for 1 h at 37°C with MNase (NEB, M0247) added in varied concentrations (17-19 Kunitz U). The reactions were quenched by adding 5 mM ethylene glycol-bis(2-aminoethylether)-N,N,N’,N’-tetraacetic acid (EGTA) (Sigma-Aldrich, E3889) and pelleted afterwards. The pellets were resuspended in PBS containing 5 mM EGTA and an aliquot of 200 mL per reaction was tested for digestion efficiency as a control. The reactions were pelleted again and resuspended in DNA ligase buffer (Thermo Scientific, B69) supplemented with dNTP mix (NEB, N0447) at a final concentration of 0.4 mM and 2.5 mM EGTA. Subsequently, 200 U/mL T4 polynucleotide Kinase (NEB, M0201), 100 U/mL DNA Polymerase I Large (Klenow) Fragment (NEB, M0210) and 300 U/mL T4 DNA ligase (Thermo Scientific, EL0013) were added. The reactions were incubated at 37°C and 20°C for 1-2 h and overnight, respectively. Following chromatin decrosslinking, DNA extraction was performed using the DNeasy blood and tissue kit (Qiagen, 69504). The MCC libraries were sonicated, indexed, and enriched with a double capture procedure as described in the Capture-C section. Biotinylated oligonucleotides (120 nt) were designed using a python-based oligo tool [64] (https://oligo.readthedocs.io/en/latest/) and ordered from IDT as xGen Lockdown Probe Pools. The final libraries were assessed on a fragment analyzer using the High Sensitivity NGS Analysis Kit and were sequenced using the NextSeq550 Illumina platform (150-bp paired-end reads). Data analysis was performed using the MCC pipeline [30].

### Tiled Micro-Capture-C

Tiled-MCC was performed using the generated MCC libraries, following a tiled enrichment procedure as described previously [14], using the Twist Hybridization and Wash Kit (Twist Bioscience, 101025). Briefly, indexed MCC libraries were pooled and dried completely in a vaccum concentrator at 45°C. Dried DNA was resuspended in blocker solution and pooled with the hybridization solution containing a custom panel of biotinylated oligonucleotides (70 nt; designed using a python-based oligo tool [64] (https://oligo.readthedocs.io/en/latest/) and incubated at 70°C overnight. Streptavidin bead pulldown and bead washes were performed with Twist Wash Buffers according to the manufacturer’s instructions (Twist Target Enrichment Protocol). Subsequently, post-hybridization PCR was performed with 11 cycles of amplification. PCR-amplified libraries were purified using pre-equilibrated Twist DNA Purification Beads. The final libraries were assessed on a fragment analyzer using the High Sensitivity NGS Analysis Kit and sequenced using the NextSeq550 Illumina platform (150-bp paired-end reads). Data analysis was performed using the MCC pipeline [30] (https://github.com/jojdavies/Micro-Capture-C) and HiC-Pro pipeline [65] (https://github.com/nservant/HiC-Pro) as described previously [14]. All contact matrices were balanced using ICE-normalization [66]. The large-scale contact matrices have a resolution of 500 bp – 2 kb (depending on the size of the region); the resolution of the nano-scale matrices is indicated in the figures.

### CUT&Tag

Cleavage under targets and tagmentation (CUT&Tag [35]) was performed for three biological replicates per experimental condition using the CUT&Tag-IT Assay Kit (Anti-Rabbit) (Active Motif, 53160) according to the manufacturer’s instructions with some modifications. Briefly, 0.5 million cells per biological replicate were mildly crosslinked with 0.3% Paraformaldehyde (Science Services, E15710) followed by quenching with 125 mM cold glycine. Meanwhile, Concanavalin A beads were prepared following the manufacturer’s instructions. The fixed cells were washed, resuspended in 1X Wash Buffer and incubated with Concanavalin A beads for 10 min on a rotator at room temperature. The tubes were placed on a magnetic stand to clear the liquid and the samples were resuspended with ice-cold Antibody Buffer supplemented with Protease Inhibitor Cocktail and Digitonin. The samples were incubated with either 1 mg SMC1A antibody (Abcam, ab9262) or 1 mg Rabbit IgG Isotype control antibody overnight at 4°C on a rotator in 0.2 mL PCR tubes. The next day, the samples were incubated with Guinea Pig Anti-Rabbit secondary antibody for 1 h at room temperature on a rotator followed by washes with Dig-Wash buffer. The tubes were placed on a magnetic stand to clear the liquid and resuspended with CUT&Tag-IT Assembled pA-Tn5 Transposomes. The reactions were subsequently incubated at room temperature on a rotator, followed by washes with Dig-300 buffer. The tubes were placed on a magnetic stand to clear the liquid and resuspended with Tagmentation Buffer. The tagmentation reactions were subsequently incubated at 37°C for 60 min. The samples were decrosslinked and DNA extraction was performed according to the manufacturer’s instructions. The purified DNA samples were PCR amplified and purified with two rounds of SPRI bead clean-up to remove primer dimers. The final libraries were assessed on a fragment analyzer using the High Sensitivity NGS Analysis Kit and sequenced using theNextSeq550 Illumina platform (75-bp paired-end reads). Paired-end reads were processed for adapter removal and duplicate filtering and mapped to the hg38 reference genome using Bowtie2 [67]. Peak calling was performed with MACS2 [68]. All peak profiles were generated using Deeptools [69].

### Public data analysis

DNase-I hypersensitivity data [70] (ENCSR000ENM) and ChIP-Seq data for CTCF [70] (ENCSR000BSE) and MED26 [24] in HCT-116 cells were analyzed using the NGseqBasic pipeline [71]. TT-seq data files for HCT-116 MED14-dTAG cells [25] were shared by the authors and differential expression analysis was performed using DESeq2 [72].

## ACKNOWLEDGEMENTS

We would like to thank Martin Jäger and Georg Winter for providing the HCT-116 MED14-dTAG cell line. We are grateful to Patrick Cramer for advice and infrastructure support. We would like to thank Michael Lidschreiber for assistance with bioinformatics analysis and Martin Jäger, Kseniia Lysakovskaia, Kerstin Maier, Petra Rus, Nadine Übelmesser, Taras Velychko and Kristina Zumer for experimental advice and support. We are also grateful to Job Dekker, Douglas Higgs, Elisa Oberbeckmann, Argyris Papantonis and Johannes Söding for helpful discussions and feedback. A.M.O. is supported by the Max Planck Society and the Deutsche Forschungsgemeinschaft.

