## Supplemental Information for "The Mediator complex regulates enhancer-promoter interactions"

**Supplementary Figure 1. Characterization of HCT-116 MED14-dTAG cells by immunoblotting and TT-seq.**

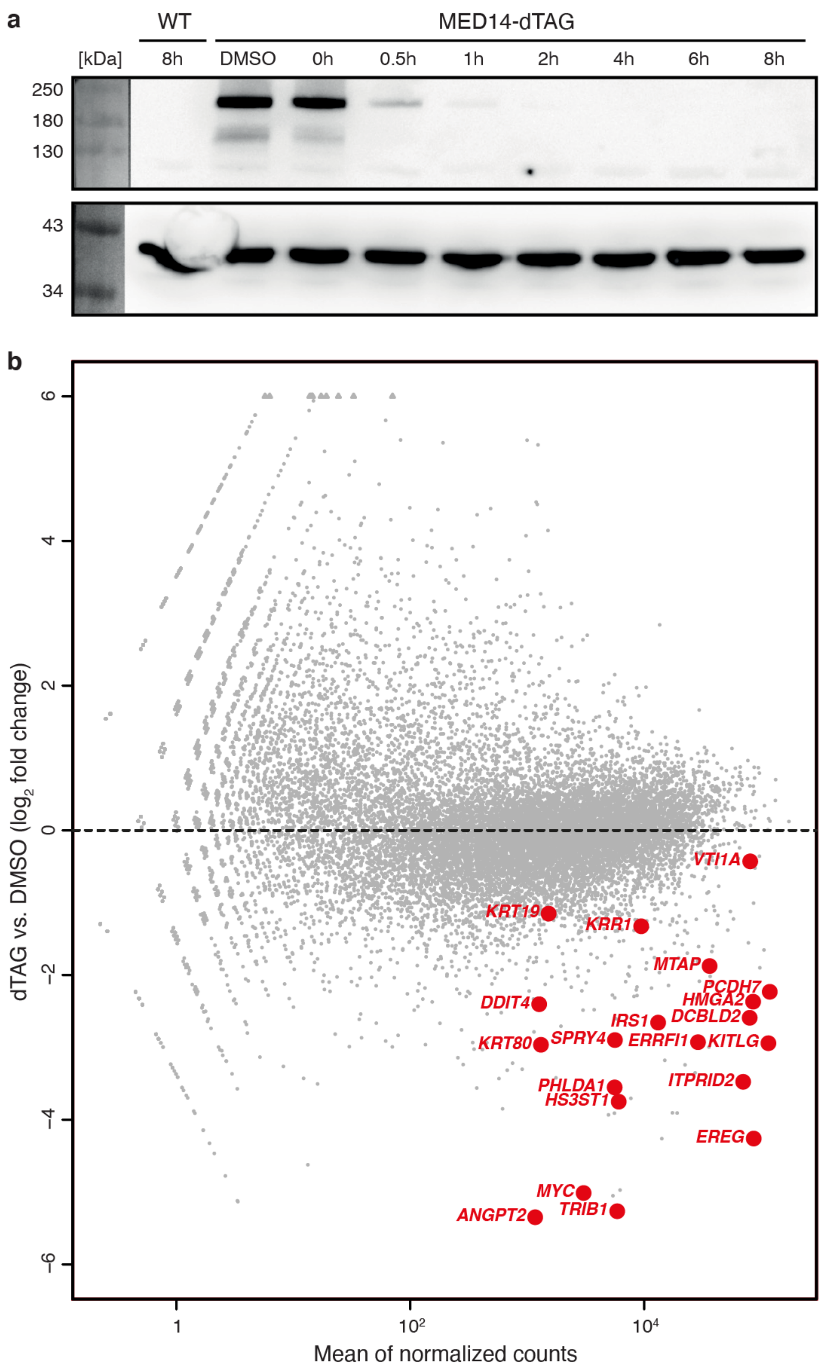


**a.** A representative immunoblot blot for MED14-dTAG-HA in wildtype (WT) and MED14-dTAG HCT-116 cells treated with DMSO for two hours or dTAG ligand for various durations as specified. MED14-dTAG-HA is detected using an anti-HA primary antibody. GAPDH is shown as a loading control. **b.** Differences in nascent transcript levels as measured by TT-seq (Jaeger et al., 2020) in HCT-116 MED14-dTAG cells treated with DMSO or dTAG ligand for two hours. Genes of interest are highlighted in red.

**Supplementary Figure 2. Capture-C analysis in HCT-116 MED14-dTAG cells.**

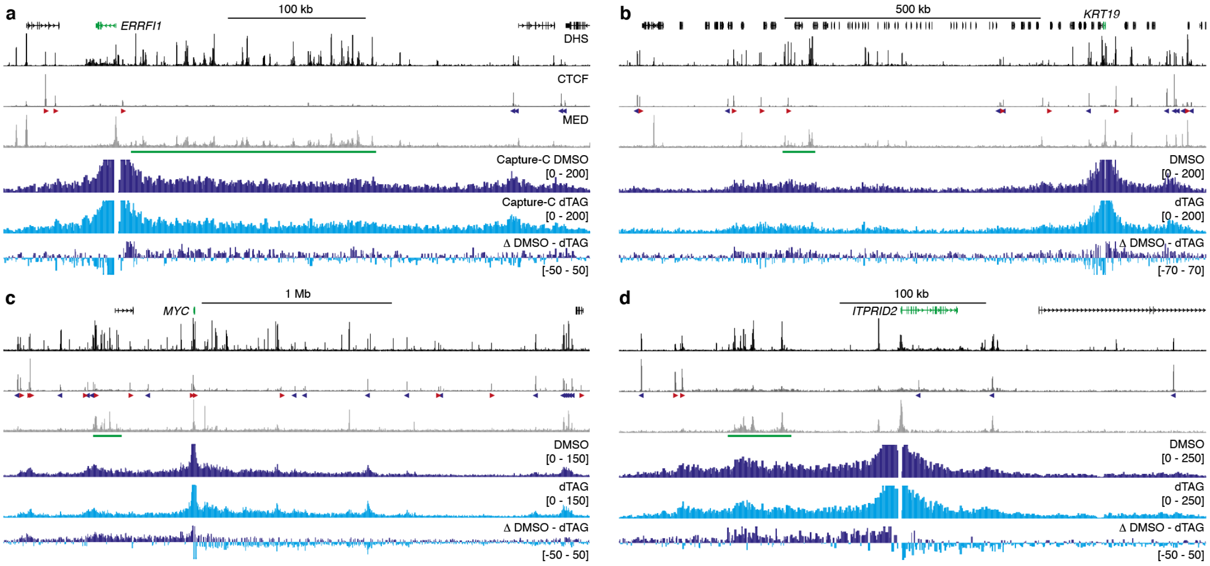


**a.** Normalized Capture-C interaction profiles from the viewpoint of the *ERRFI1* promoter in HCT-116 MED14-dTAG cells treated for two hours with DMSO (dark blue) or dTAG ligand (light blue). Gene annotation, DNase hypersensitive sites (DHS), and ChIP-seq data for CTCF and MED26 are shown above. Enhancers of interest are highlighted in green below the MED26 profiles and orientations of CTCF motifs are indicated with arrowheads (forward orientation in red; reverse orientation in blue). The axes of the DHS and ChIP-seq profiles are scaled to signal and have the following ranges: DHS = 0–170; CTCF = 0–253; MED26 = 0–56. Coordinates (hg38): chr1:7,945,000-8,370,000. **b.** Data as described in **a** for the *KRT19* locus. The axes of the DHS and ChIP-seq profiles are scaled to signal and have the following ranges: DHS = 0–180; CTCF = 0–301; MED26 = 0–77. Coordinates (hg38): chr17:40,580,000-41,725,000. **c.** Data as described in **a** for the *MYC* locus. The axes of the DHS and ChIP-seq profiles are scaled to signal and have the following ranges: DHS = 0–188; CTCF = 0–188; MED26 = 0–64. Coordinates (hg38): chr8:126,735,000-129,820,000. **d.** Data as described in **a** for the *ITPRID2* locus. The axes of the DHS and ChIP-seq profiles are scaled to signal and have the following ranges: DHS = 0–173; CTCF = 0–69; MED26 = 0–68. Coordinates (hg38): chr2:181,700,000-182,100,000.

**Supplementary Figure 3. Micro-Capture-C analysis in HCT-116 MED14-dTAG cells.**

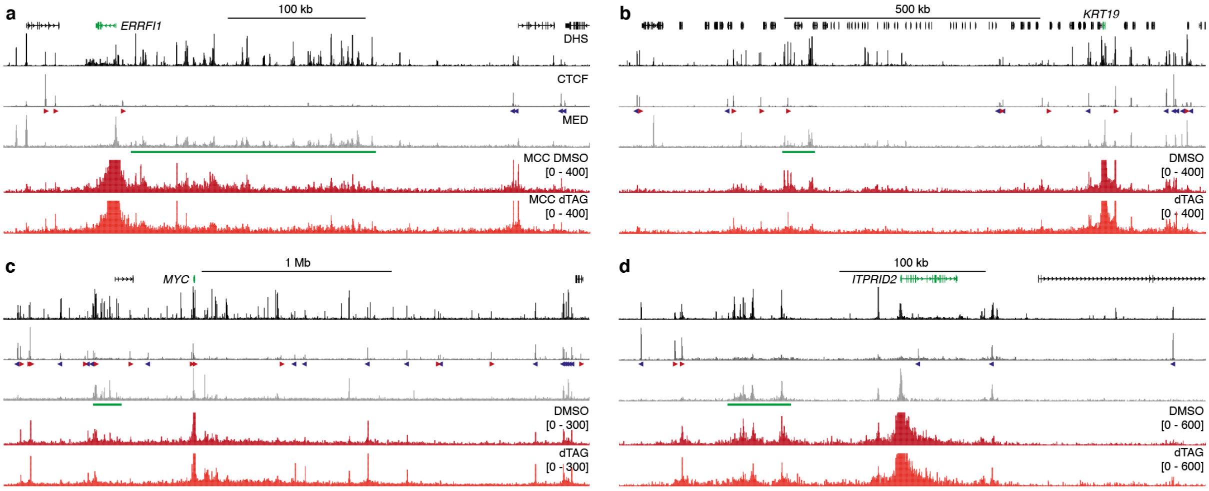


**a.** Normalized Micro-Capture-C interaction profiles from the viewpoint of the *ERRFI1* promoter in HCT-116 MED14-dTAG cells treated for two hours with DMSO (dark red) or dTAG ligand (light red). Gene annotation, DNase hypersensitive sites (DHS), and ChIP-seq data for CTCF and MED26 are shown above. Enhancers of interest are highlighted in green below the MED26 profiles and orientations of CTCF motifs are indicated with arrowheads (forward orientation in red; reverse orientation in blue). The axes of the DHS and ChIP-seq profiles are scaled to signal and have the following ranges: DHS = 0–170; CTCF = 0–253; MED26 = 0–56. Coordinates (hg38): chr1:7,945,000-8,370,000. **b.** Data as described in **a** for the *KRT19* locus. The axes of the DHS and ChIP-seq profiles are scaled to signal and have the following ranges: DHS = 0–180; CTCF = 0–301; MED26 = 0–77. Coordinates (hg38): chr17:40,580,000-41,725,000. **c.** Data as described in **a** for the *MYC* locus. The axes of the DHS and ChIP-seq profiles are scaled to signal and have the following ranges: DHS = 0–188; CTCF = 0–188; MED26 = 0–64. Coordinates (hg38): chr8:126,735,000-129,820,000. **d.** Data as described in **a** for the *ITPRID2* locus. The axes of the DHS and ChIP-seq profiles are scaled to signal and have the following ranges: DHS = 0–173; CTCF = 0–69; MED26 = 0–68. Coordinates (hg38): chr2:181,700,000-182,100,000.

**Supplementary Figure 4. Tiled-MCC analysis of the *MTAP* locus in HCT-116 MED14-dTAG cells.**

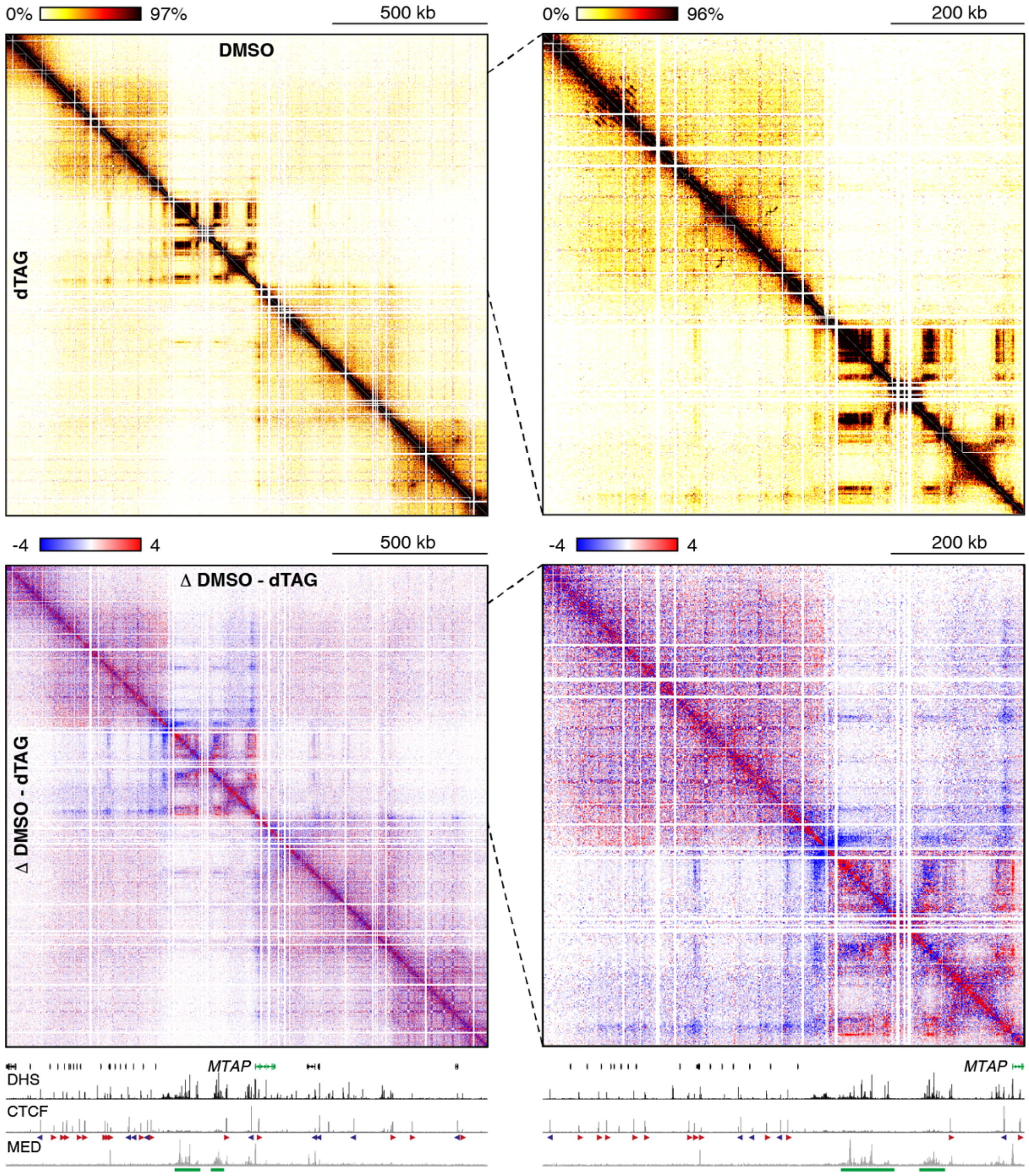


Tiled-MCC contact matrices of the *MTAP* locus in HCT-116 MED14-dTAG cells treated for two hours with DMSO (top-right) or dTAG ligand (bottom-left). The right matrix shows a zoomed view, of which the area is indicated by the black dashed lines. Differential contact matrices, in which interactions enriched in DMSO-treated cells are shown in red and interactions enriched in dTAG-treated cells are shown in blue, are displayed below. Gene annotation, DNase hypersensitive sites (DHS), and ChIP-seq data for CTCF and MED26 are shown at the bottom. Enhancers of interest are highlighted in green below the MED26 profiles and orientations of CTCF motifs are indicated with arrowheads (forward orientation in red; reverse orientation in blue). The axes of the DHS and ChIP-seq profiles are scaled to signal and have the following ranges: DHS = 0–186; CTCF = 0–112; MED26 = 0–62. Coordinates (hg38): chr9:21,000,000-22,550,000.

**Supplementary Figure 5. Tiled-MCC analysis of the *HMGA2* locus in HCT-116 MED14-dTAG cells.**

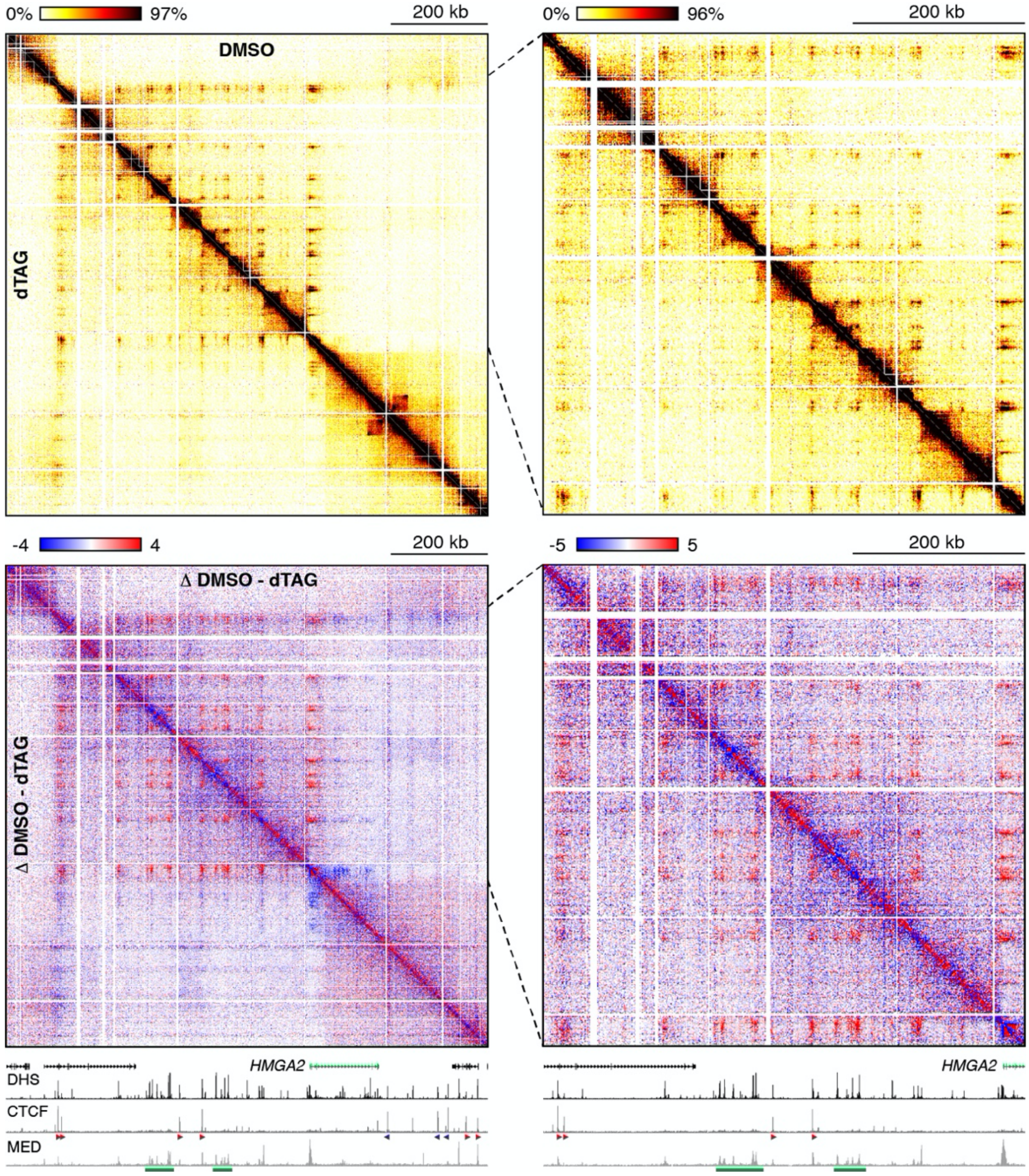


Tiled-MCC contact matrices of the *HMGA2* locus in HCT-116 MED14-dTAG cells treated for two hours with DMSO (top-right) or dTAG ligand (bottom-left). The right matrix shows a zoomed view, of which the area is indicated by the black dashed lines. Differential contact matrices, in which interactions enriched in DMSO-treated cells are shown in red and interactions enriched in dTAG-treated cells are shown in blue, are displayed below. Gene annotation, DNase hypersensitive sites (DHS), and ChIP-seq data for CTCF and MED26 are shown at the bottom. Enhancers of interest are highlighted in green below the MED26 profiles and orientations of CTCF motifs are indicated with arrowheads (forward orientation in red; reverse orientation in blue). The axes of the DHS and ChIP-seq profiles are scaled to signal and have the following ranges: DHS = 0–181; CTCF = 0–93; MED26 = 0–66. Coordinates (hg38): chr12:65,200,000-66,190,000.

**Supplementary Figure 6. Tiled-MCC analysis of the *ITPRID2* locus in HCT-116 MED14-dTAG cells.**

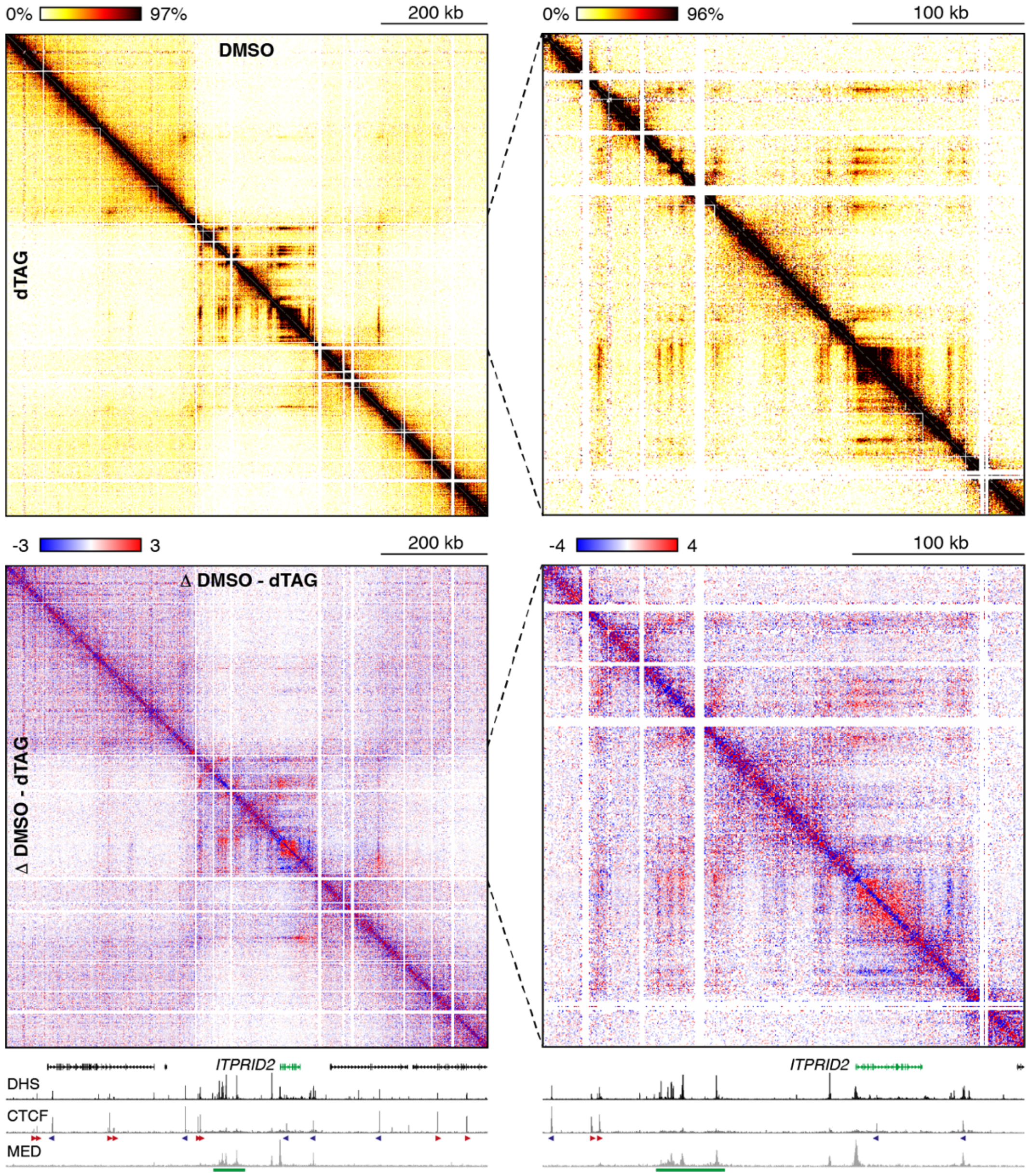


Tiled-MCC contact matrices of the *ITPRID2* locus in HCT-116 MED14-dTAG cells treated for two hours with DMSO (top-right) or dTAG ligand (bottom-left). The right matrix shows a zoomed view, of which the area is indicated by the black dashed lines. Differential contact matrices, in which interactions enriched in DMSO-treated cells are shown in red and interactions enriched in dTAG-treated cells are shown in blue, are displayed below. Gene annotation, DNase hypersensitive sites (DHS), and ChIP-seq data for CTCF and MED26 are shown at the bottom. Enhancers of interest are highlighted in green below the MED26 profiles and orientations of CTCF motifs are indicated with arrowheads (forward orientation in red; reverse orientation in blue). The axes of the DHS and ChIP-seq profiles are scaled to signal and have the following ranges: DHS = 0–173; CTCF = 0–69; MED26 = 0–68. Coordinates (hg38): chr2:181,380,000-182,280,000.

**Supplementary Figure 7. SMC1A CUT&Tag analysis in HCT-116 MED14-dTAG cells.**

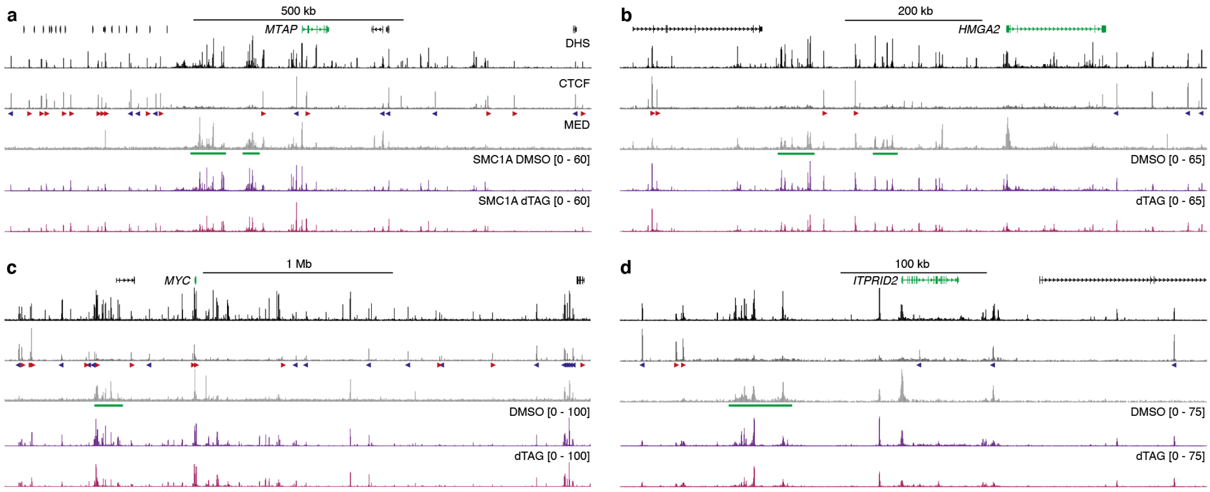


**a.** CUT&Tag data for the Cohesin subunit SMC1A in the *MTAP* locus in HCT-116 MED14-dTAG cells treated for two hours with DMSO (dark purple) or dTAG ligand (light purple). Gene annotation, DNase hypersensitive sites (DHS), and ChIP-seq data for CTCF and MED26 are shown above. Enhancers of interest are highlighted in green below the MED26 profiles and orientations of CTCF motifs are indicated with arrowheads (forward orientation in red; reverse orientation in blue). The axes of the DHS and ChIP-seq profiles are scaled to signal and have the following ranges: DHS = 0–186; CTCF = 0–112; MED26 = 0–62. Coordinates (hg38): chr9:21,096,000-22,491,000. **b.** Data as described in **a** for the *HMGA2* locus. The axes of the DHS and ChIP-seq profiles are scaled to signal and have the following ranges: DHS = 0–181; CTCF = 0–93; MED26 = 0–66. Coordinates (hg38): chr12:65,260,000-66,115,000. **c.** Data as described in **a** for the *MYC* locus. The axes of the DHS and ChIP-seq profiles are scaled to signal and have the following ranges: DHS = 0–188; CTCF = 0–188; MED26 = 0–64. Coordinates (hg38): chr8:126,735,000-129,820,000. **d.** Data as described in **a** for the *ITPRID2* locus. The axes of the DHS and ChIP-seq profiles are scaled to signal and have the following ranges: DHS = 0–173; CTCF = 0–69; MED26 = 0–68. Coordinates (hg38): chr2:181,700,000-182,100,000.

**Supplementary Figure 8. Micro-topology analysis of the *MTAP* locus in HCT-116 MED14-dTAG cells.**

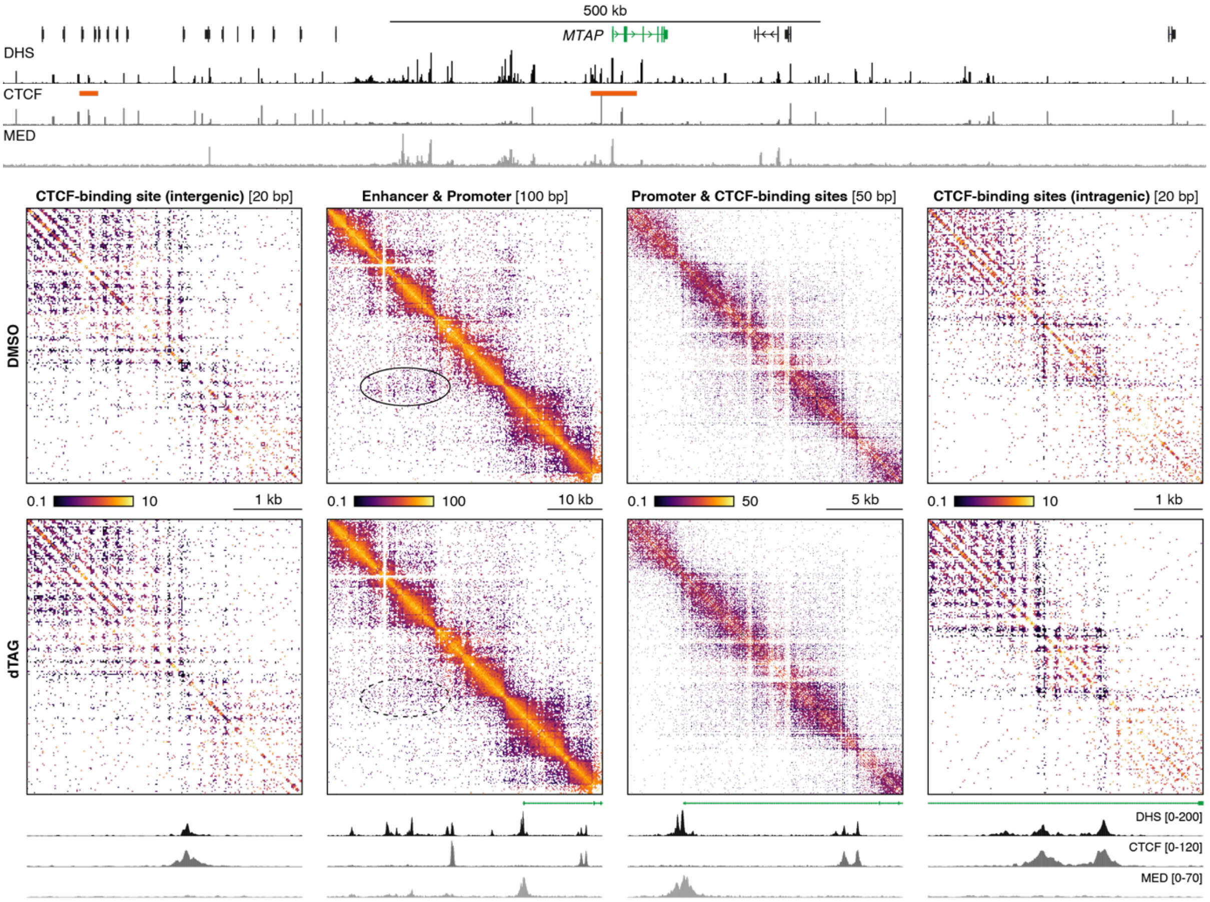


Tiled-MCC ligation junctions in the *MTAP* locus in HCT-116 MED14-dTAG cells treated for two hours with DMSO (top) or dTAG ligand (bottom), displayed in localized contact matrices at high resolution. Gene annotation, DNase hypersensitive sites (DHS), and ChIP-seq data for CTCF and MED26 for the extended and localized *MTAP* locus are shown above and below the matrices, respectively. The regions covered in the contact matrices are highlighted with orange bars (not drawn to scale) below the top DHS profile and show an intergenic CTCF-binding site, an interacting enhancer and promoter, a promoter and CTCF-binding sites, and intragenic CTCF-binding sites, at the indicated resolution. Note that the three matrices on the right show partially overlapping regions. The axes of the DHS and ChIP-seq profiles at the bottom are fixed as indicated; the profiles at the top are scaled to signal with the following ranges: DHS = 0–186; CTCF = 0–112; MED26 = 0–62.

**Supplementary Figure 9. Micro-topology analysis of the *HMGA2* locus in HCT-116 MED14-dTAG cells.**

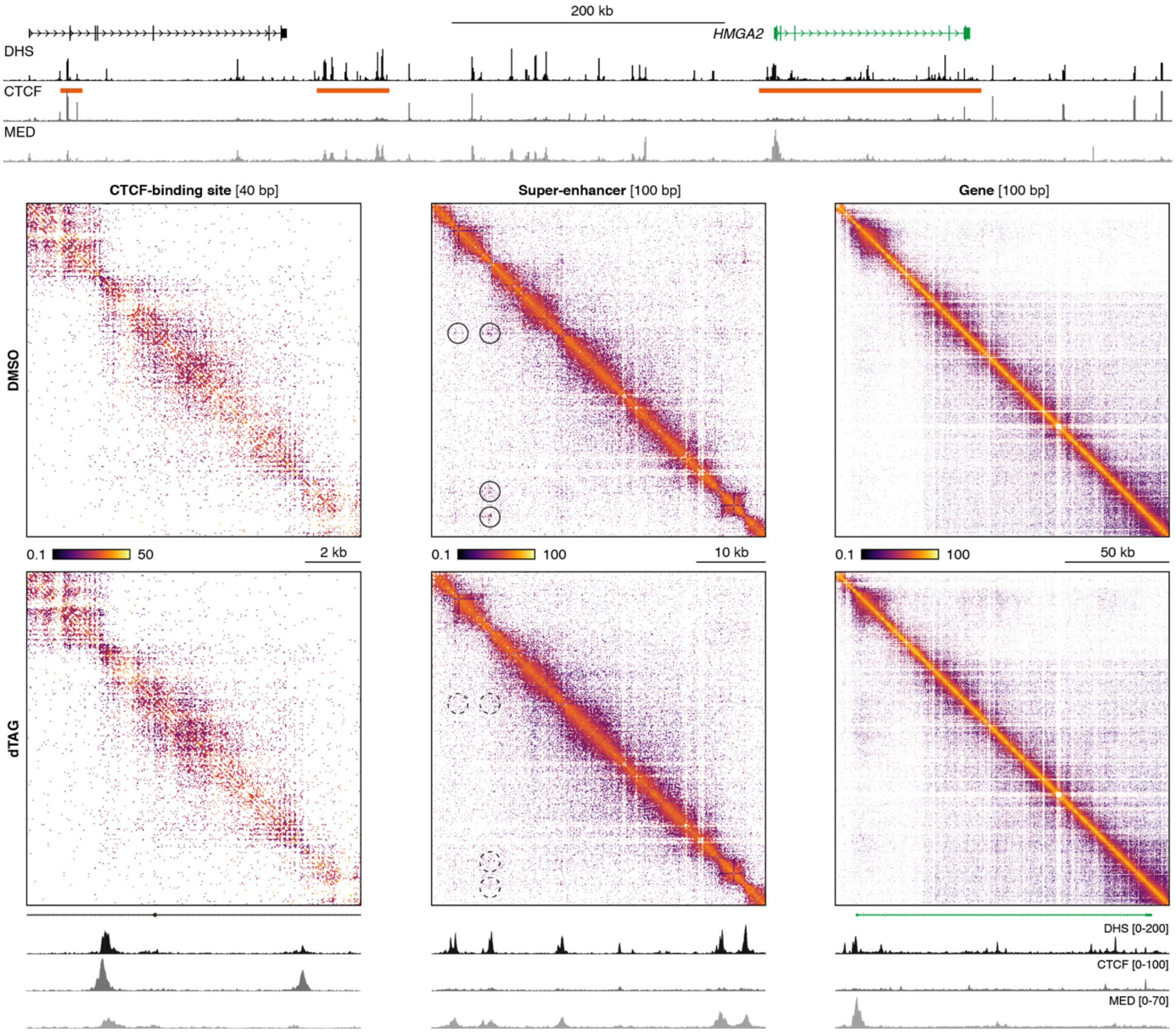


Tiled-MCC ligation junctions in the *HMGA2* locus in HCT-116 MED14-dTAG cells treated for two hours with DMSO (top) or dTAG ligand (bottom), displayed in localized contact matrices at high resolution. Gene annotation, DNase hypersensitive sites (DHS), and ChIP-seq data for CTCF and MED26 for the extended and localized *HMGA2* locus are shown above and below the matrices, respectively. The regions covered in the contact matrices are highlighted with orange bars (not drawn to scale) below the top DHS profile and show a CTCF-binding site, a super-enhancer, and a gene, at the indicated resolution. The axes of the DHS and ChIP-seq profiles at the bottom are fixed as indicated; the profiles at the top are scaled to signal with the following ranges: DHS = 0–181; CTCF = 0–93; MED26 = 0–66.

**Supplementary Figure 10. Micro-topology analysis of the *ITPRID2* locus in HCT-116 MED14-dTAG cells.**

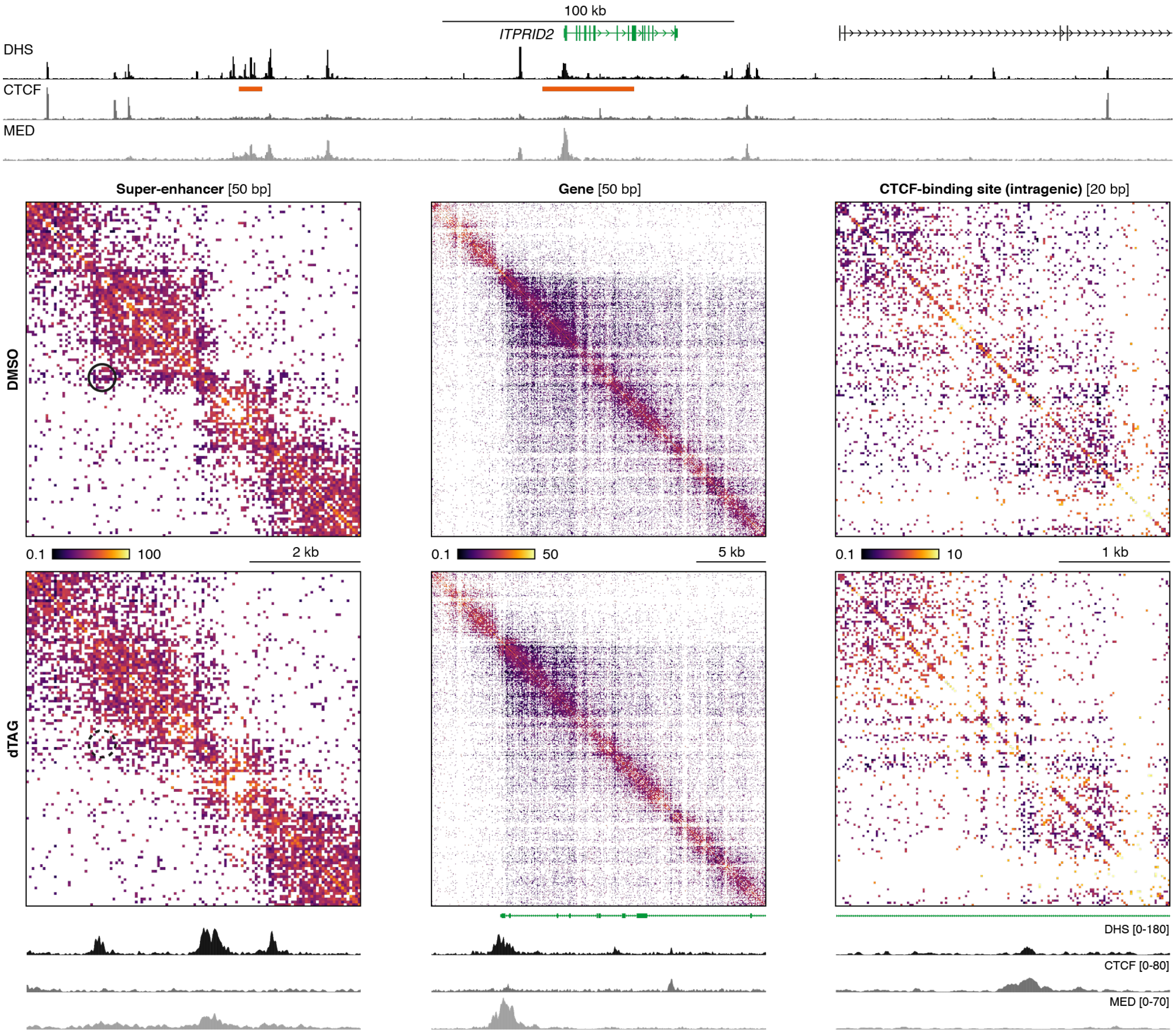


Tiled-MCC ligation junctions in the *ITPRID2* locus in HCT-116 MED14-dTAG cells treated for two hours with DMSO (top) or dTAG ligand (bottom), displayed in localized contact matrices at high resolution. Gene annotation, DNase hypersensitive sites (DHS), and ChIP-seq data for CTCF and MED26 for the extended and localized *ITPRID2* locus are shown above and below the matrices, respectively. The regions covered in the contact matrices are highlighted with orange bars (not drawn to scale) below the top DHS profile and show a super-enhancer, a gene, and an intragenic CTCF-binding site, at the indicated resolution. The axes of the DHS and ChIP-seq profiles at the bottom are fixed as indicated; the profiles at the top are scaled to signal with the following ranges: DHS = 0–173; CTCF = 0–69; MED26 = 0–68.

**Supplementary Figure 11. Combined MED14 depletion and BET inhibition does not lead to further weakening of enhancer-promoter interactions.**

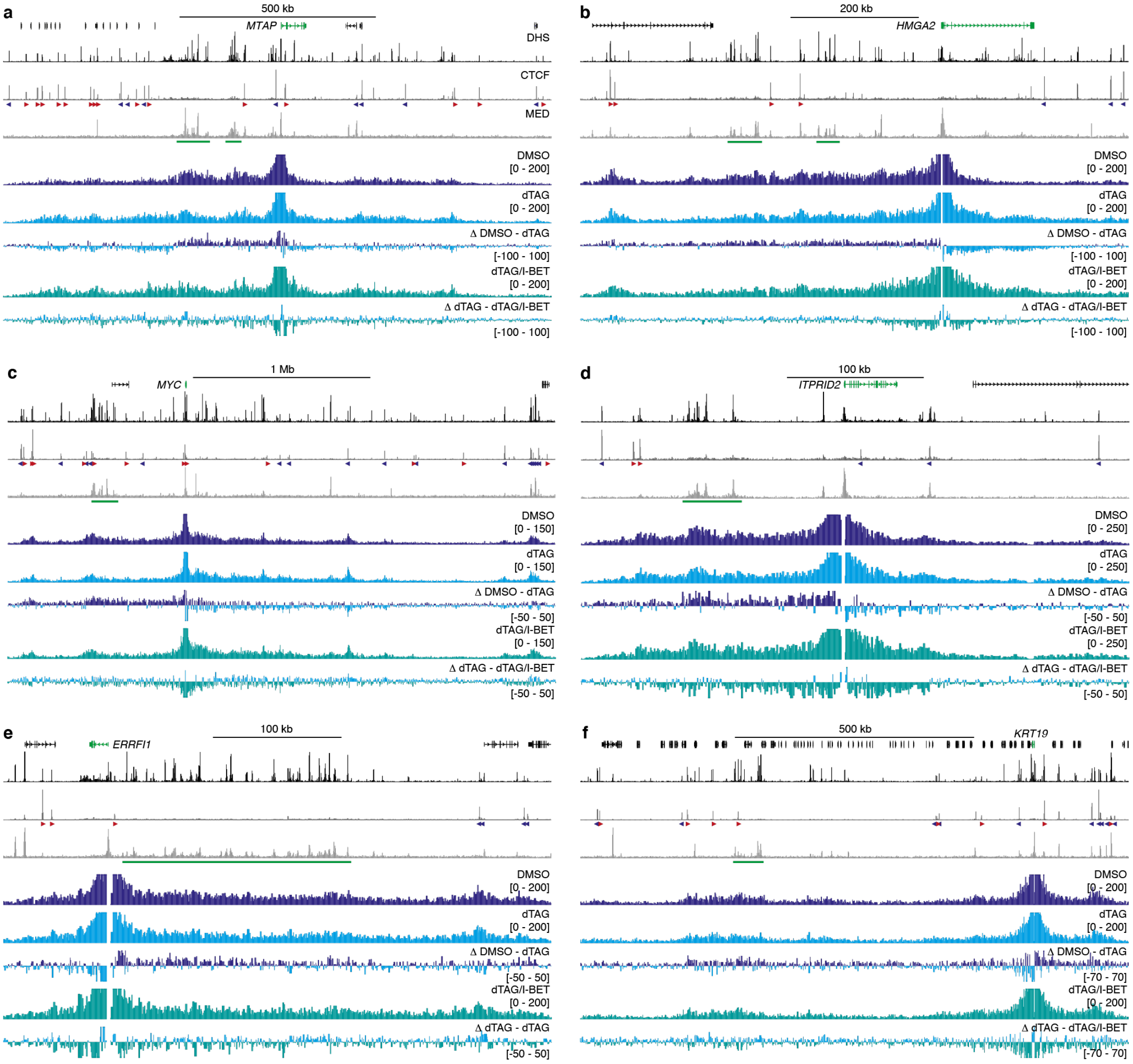


**a.** Normalized Capture-C interaction profiles from the viewpoint of the *MTAP* promoter in HCT-116 MED14-dTAG cells for the following conditions: (I) treatment with DMSO for two hours (dark blue); (II) treatment with dTAG ligand for two hours (light blue); treatment with dTAG ligand for two hours and I-BET for 90 minutes (teal). Gene annotation, DNase hypersensitive sites (DHS), and ChIP-seq data for CTCF and MED26 are shown above. Enhancers of interest are highlighted in green below the MED26 profiles and orientations of CTCF motifs are indicated with arrowheads (forward orientation in red; reverse orientation in blue). The axes of the DHS and ChIP-seq profiles are scaled to signal and have the following ranges: DHS = 0–186; CTCF = 0–112; MED26 = 0–62. Coordinates (hg38): chr9:21,096,000-22,491,000. **b.** Data as described in **a** for the *HMGA2* locus. The axes of the DHS and ChIP-seq profiles are scaled to signal and have the following ranges: DHS = 0–181; CTCF = 0–93; MED26 = 0–66. Coordinates (hg38): chr12:65,260,000-66,115,000. **c.** Data as described in **a** for the *MYC* locus. The axes of the DHS and ChIP-seq profiles are scaled to signal and have the following ranges: DHS = 0–188; CTCF = 0–188; MED26 = 0–64. Coordinates (hg38): chr8:126,735,000-129,820,000. **d.** Data as described in **a** for the *ITPRID2* locus. The axes of the DHS and ChIP-seq profiles are scaled to signal and have the following ranges: DHS = 0–173; CTCF = 0–69; MED26 = 0–68. Coordinates (hg38): chr2:181,700,000-182,100,000. **e.** Data as described in **a** for the *ERRFI1* locus. The axes of the DHS and ChIP-seq profiles are scaled to signal and have the following ranges: DHS = 0–170; CTCF = 0–253; MED26 = 0–56. Coordinates (hg38): chr1:7,945,000-8,370,000. **f.** Data as described in **a** for the *KRT19* locus. The axes of the DHS and ChIP-seq profiles are scaled to signal and have the following ranges: DHS = 0–180; CTCF = 0–301; MED26 = 0–77. Coordinates (hg38): chr17:40,580,000-41,725,000
